# Inferring Gene Regulatory Networks from Single Cell RNA-seq Temporal Snapshot Data Requires Higher Order Moments

**DOI:** 10.1101/2021.05.05.440762

**Authors:** N. Alexia Raharinirina, Felix Peppert, Max von Kleist, Christof Schütte, Vikram Sunkara

## Abstract

Single cell RNA-sequencing (scRNA-seq) has become ubiquitous in biology. Recently, there has been a push for using scRNA-seq snapshot data to infer the underlying gene regulatory networks (GRNs) steering cellular function. To date, this aspiration remains unrealised due to technical- and computational challenges. In this work, we focus on the latter, which is under-represented in the literature.

We took a systemic approach by subdividing the GRN inference into three fundamental components: the data pre-processing, the feature extraction, and the inference. We saw that the regulatory signature is captured in the statistical moments of scRNA-seq data, and requires computationally intensive minimisation solvers to extract. Furthermore, current data pre-processing might not conserve these statistical moments.

Though our moment-based approach is a didactic tool for understanding the different compartments of GRN inference, this line of thinking–finding computationally feasible multi-dimensional statistics of data–is imperative for designing GRN inference methods.

## 1 Introduction

The emergence of single cell RNA sequencing technology (scRNA-seq), the extraction of the transcriptome of individual cells, has helped immensely in detecting and delineating heterogeneities in cells (1; 2; 3; 4; 5; 6; 7). Furthermore, with advances in machine learning and mRNA metabolic tagging, scRNA-seq has given new insights into cellular development and disease pathogenesis (2; 5; 6; 7; 8; 9; 10). In light of these advances, the development of methods which infer the underlying *gene regulatory network* (GRN)–which drives cellular decisions–is lagging behind (11).

Cellular function is dependent on the cell’s transcriptomic signature, where the proteins translated from the mRNA form signalling pathways, which perform the cellular function and then in a feedback loop, regulate the mRNA transcription to then translate proteins (12; 13; 14). The process of a gene affecting the expression of another gene is referred to as *gene regulation*, and the collection of all gene regulatory interactions (e.g., in a cell) forms a gene regulatory network (GRN) (3; 7; 13; 15; 16; 17; 18). Unlike protein-protein interactions, where educts are converted into products, gene regulatory interactions are more illusive. A gene regulates another gene through its downstream protein complexes, which affect the rate of transcription of the gene being regulated. That is, gene regulation physically occurs on the DNA level and its effect is observed on the mRNA level. In particular, “Gene A regulates Gene B” means that gene A either up-regulates (promotes) or down-regulates (inhibits) the rate of transcription of gene B. The fact that scRNA-seq only captures mRNA, while gene regulation interactions take place up- or down stream from the mRNA, constitutes a major hurdle for inferring GRNs from scRNA-seq data (19; 20). Recently, a trend has emerged to use multiple temporal scRNA-seq snapshots to capture the underlying GRNs of cells.

Current single-cell GRN inference methods using temporal snapshot data can be grouped into two families: the distribution-based methods and the moment-based methods, distinguished by the type of summary statistics that they use. Distribution-based methods construct their summary statistic based on the empirical distribution of the gene expression in the snapshots (21; 22; 23; 24; 25), whereas, moment-based methods utilise only the moments (mean and covariance) of the snapshot data (26; 27; 28).

The rapid increase in the number of GRN inference methods has motivated the development of comprehensive comparative frameworks. A recent benchmark of twelve GRN methods demonstrated that the algorithms struggled to predict the ground truth GRNs, and speculated that the low performance was due to the insufficient resolution in the scRNA-seq data (11). Rather than proposing another method, the focus of this paper is to dissect and identify key computational stumbling blocks for inferring GRNs from scRNA-seq data.

In general, there are three key components to a GRN inference method: the data *pre-processing*, the *feature extraction*, and the *inference* of the underlying gene regulation pattern. The data pre-processing transforms the data to make it more tractable for analysis, for example, pseudotime reordering (29) and variational autoen-coders (30). The feature extraction compiles summary statistic of the data which is intended to contain the information of the regulation, for example, the mean or mutual information. Lastly, the inference of the underlying gene regulation pattern is what finds the GRN among the space of all possible GRNs which best matches the statistics of the data, for example, linear least squares or random forest. We take a bottom-up approach and shed light on key challenges in these three core components through intuitive tailored GRN models.

In this work, we construct three different in-silico GRN (known ground truth), containing different gene regulation patterns, and modelled by Markov-jump processes according to the standard dogma, with the regulation placed up-stream from the mRNA (14; 31). Our GRN models consider the intrinsic noise arising from the stochastic nature of gene regulation (32; 33; 34; 35). We designed the models to be nested with ascending order in the number of regulatory reactions, from no regulatory interactions (only correlation) to many regulatory interactions (double feed-back loop). We then use simulations to produce artificial/synthetic scRNA-seq data on different levels of resolution (e.g. time lags). We omit extrinsic noise in order to investigate, in the best case scenario, the possibility of reconstructing the three GRN models by four inference methods. Our strategy is based on the reasoning that if a GRN inference method is not able to infer the ground truth for clean artificial data, then it won’t be successful for real-world data which contain a multitude of caveats. For comparison, we chose two distribution-based inference methods: mutual information (22) and SINCERITIES (24), and two moment-based methods: moments derivative-based (Linear moment-based inference (26)) and higher-order moment-based (Non-linear moment-based inference [this paper]).

We finish this work by discussing the need for higher-order moments for GRN inference and their computational challenges. We will also discuss the caveats of existing datasets and the need for multi-omics data for truly validating GRNs inferred from scRNA-seq temporal snapshot data.

## 2 Results

### 2.1 The variance time course is only moderately explained by the mean time course in single cell datasets

Most GRN inference methods use the mean time course to infer their GRNs. This is historically motivated by bulk RNA-seq, where only the mean transcription levels are observed. However, with single cell transcriptomics, more complex statistical time courses can be constructed. To illustrate the information captured by more complex statistics, we studied the trends in the mean- and variance of gene expression of individual genes from single cell datasets. We used the variance as a simple surrogate of a non-linear statistic and quantified the correspondence between trends in the mean and the trends in the variance across five different single cell datasets (see Methods 4.1).

Looking at the hESC scRNA-seq dataset, we saw that the trends of the mean time course and the trends of the variance time course followed similar behaviour (Fig. 1 a). However, when we looked at the correspondence of individual genes, we found that genes which had a similar mean trend exhibited different variance trends, for example, genes following mean trend number 4 (a trend of: up, flat, then down) were distributed mostly across trends a, b, and d in the variance (Fig. 1b). Similar patterns were observed for the other datasets. We further quantified this correspondence using the measure of *proficiency* (also known as normalised mutual information) to compare between the datasets (Fig. 1 c). We found that real datasets had proficiency in the range of 6 % to 37 %. That is, 6 to 37 percent of the trends in the variance time courses corresponded with the trends of the mean time courses. Furthermore, we found that the synthetic dataset made through GeneNetWeaver (GNW) had very low proficiency (below 5 %), which could be attributed to the use of white noise in its model. In contrast, the synthetic dataset from the BoolODE method had a high proficiency (above 50 %), which could be attributed to the use of mean-based coloured noise in its model.

**Figure 1:**
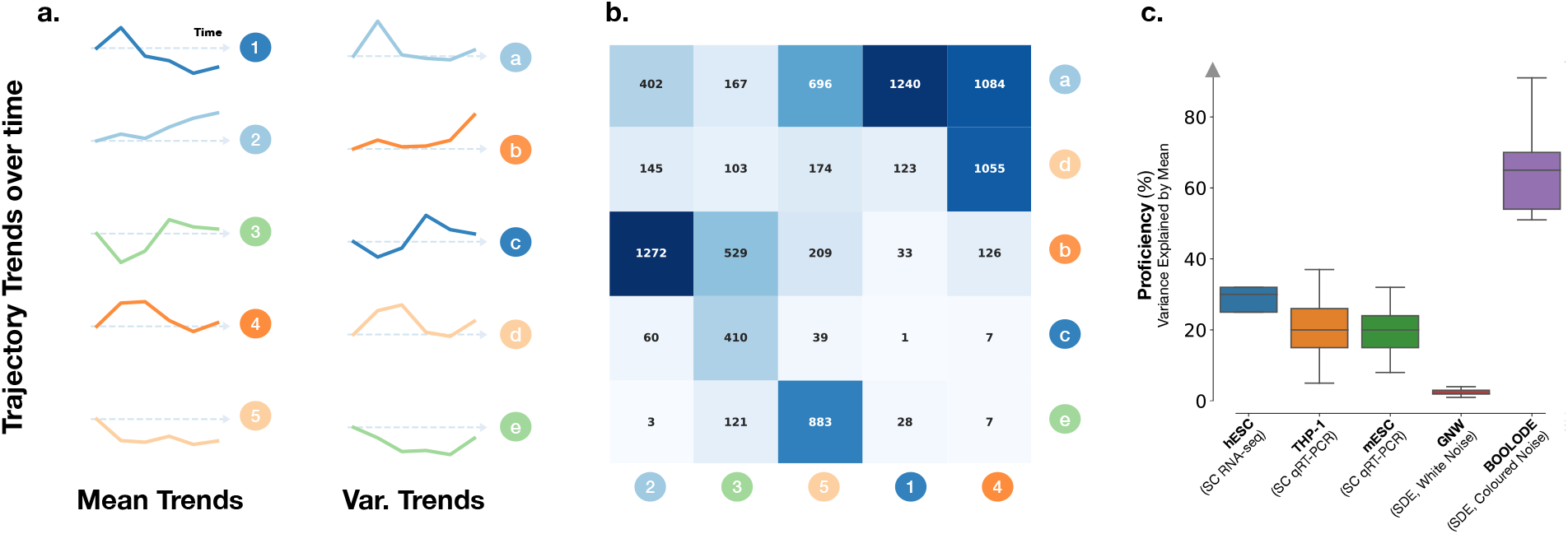
Mean trends and variance trends in single cell datasets. **a.** Trends in the mean- and variance of gene expression for hESC scRNA-seq dataset. **b.** The number of genes in the hESC scRNA-seq dataset associated with the mean and variance trends. **c.** Percentage of variance trends explained by mean trends across five different single cell datasets.

In summary, the mean- and the variance time courses of real gene expression data contained mutually exclusive information. This gives motivation to enrich GRN inference through the use of more complex statistics, for example, the higher order moments.

### 2.2 Stochastic Two Gene Interaction Models

To investigate the role of moments in unravelling regulatory reactions from scRNA-seq data, we constructed three simple two-gene GRN models (see Methods 4.2). Our first GRN model was a simple no interaction (No-I) two gene model, where each gene, Gene A and Gene B, can be in one of two discrete states, on or basal (off state), and can switch between these states via a constant propensity. The gene is then transcribed into mRNA at a constant rate depending on the state of the gene. The transcribed mRNA then undergoes translation and the respective protein is synthesised (Fig. 2 Top). The mRNA and proteins undergo degradation proportional to their respective populations. In the No-I model, the downstream products associated to their respective gene are not correlated across genes. Our second GRN model was a mono-directional interaction (Mono-I) model, that is, it had the same reactions as the No-I model with the exception of an interaction where Protein B actively upregulates the switching off of Gene A (Fig. 2 Middle). In this scenario, Gene A and its downstream products are affected by the regulation of Gene B, however, Gene B is not affected by any downstream products of Gene A. Lastly, our third GRN model was the bi-directional interaction (Bi-I) model, where protein A upregulates the switching off of Gene B and vice versa, protein B upregulates the switching off of Gene A (Fig. 2 Bottom). In the Bi-I model, all products in the system are correlated.

**Figure 2:**
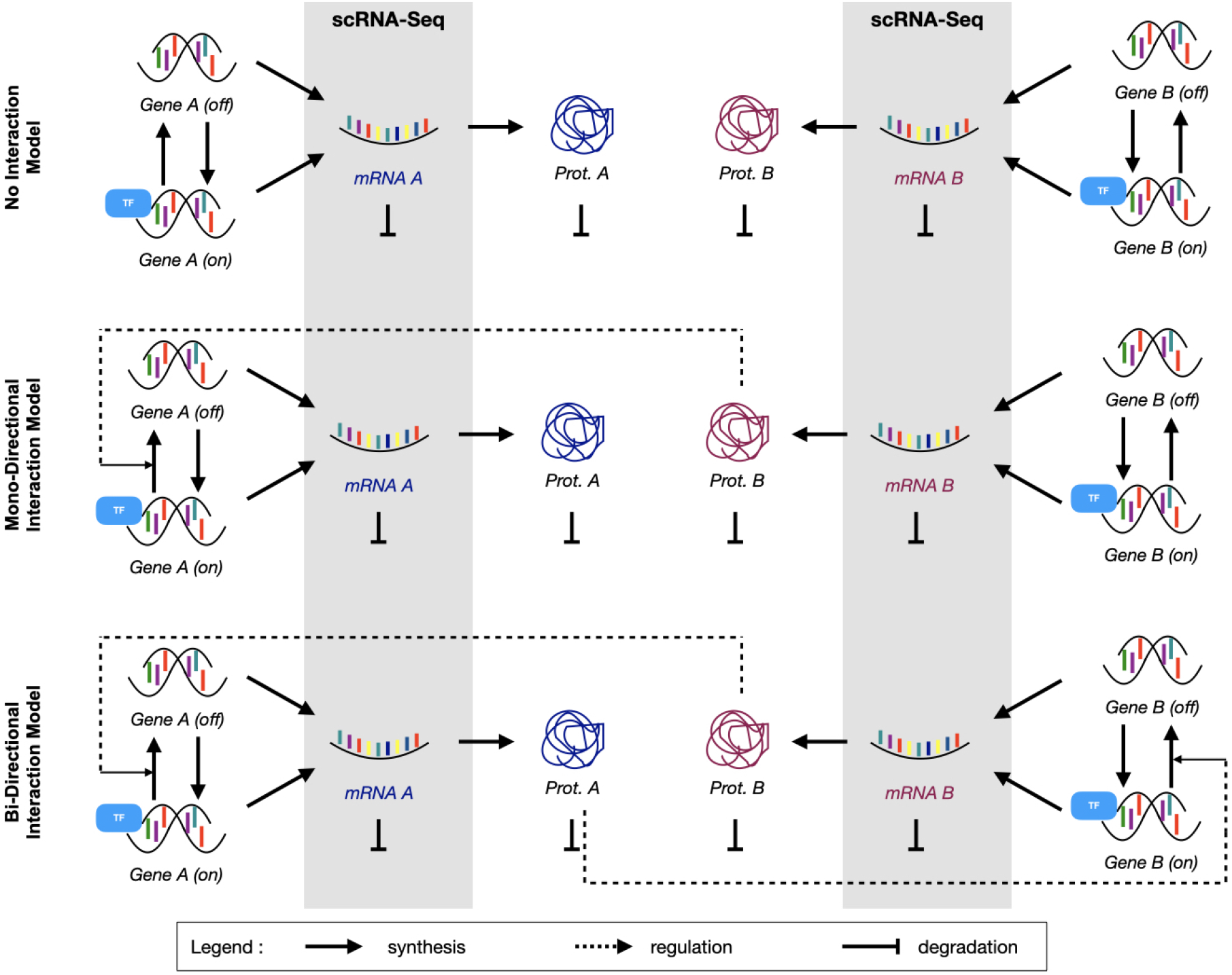
Two gene interaction models. The three GRN models of interest comprising two Genes A and B and their corresponding mRNA and proteins. From top to bottom, model schematics for **No-I:** no interaction model, **Mono-I:** mono-directional interaction, and **Bi-I:** bi-directional interaction. Only the mRNA counts, shown in the grey shaded area, are used in the GRN inference methods.

#### 2.2.1 Covariance and Skewness can aid in Detecting Regulatory Pathways

The three models were simulated using the Stochastic Simulation Algorithm (SSA) (see Methods 4.3). Only the mRNA expression counts from the simulations were extracted for regulatory inference, to mimic scRNA-seq data.

In the No-I model, we observed that both the time course of the mean expression of both mRNAs (A and B) increased identically until the time horizon (Fig. 3 a, Sup. Fig. A a). Due to there being no interactions across genes, as expected, the samples at any fixed time point exhibited near zero covariance between the mRNA expression counts (Fig. 3 a,e). In the Mono-I model, we observed that at early time points, the mRNAs’ mean expression increased similarly, then, the mean expression of mRNA A started to plateau while the mean expression of mRNA B continued to rise, and had a similar time course as the mRNA B in the No-I model (Fig. 3 b, Sup. Fig. A b). We observed in the time course, that the mRNAs had a negative covariance between them (Fig. 3 b,e). Lastly, in the Bi-I model, we observed that the mean expression time course of the mRNAs increased identically–like in the No-I model. The expression distribution was found to also have a negative covariance structure, however, upon inspecting the distribution of a snapshot at *T* = 60 min, we saw that the distribution was very symmetric and was shaped like a waning crescent (Fig. 3 c-e).

**Figure 3:**
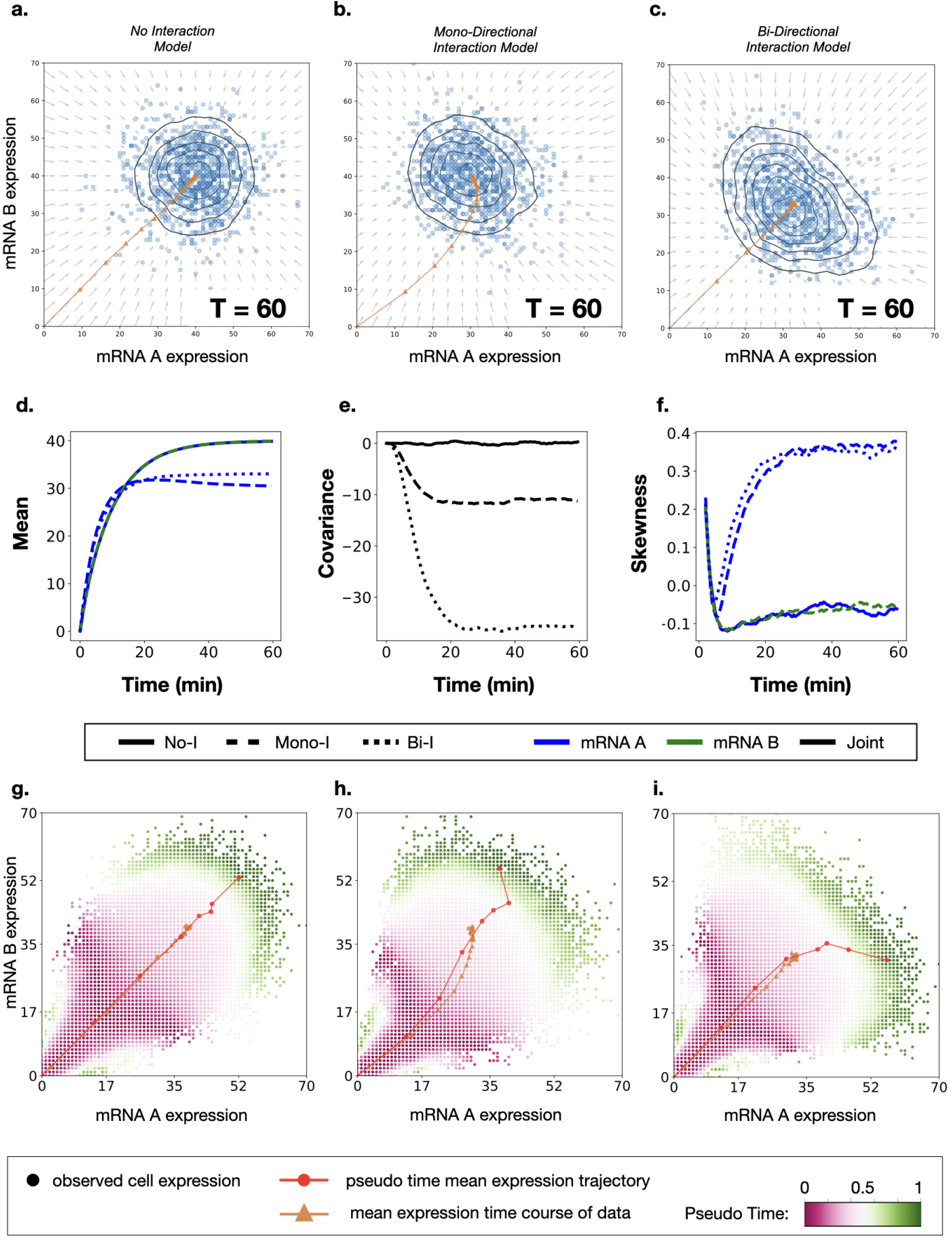
Snapshots of the three interaction models. Snapshot gene expression data at time *T* = 60 showing 1000 sample mRNA population counts. **a.** to **c.** correspond to No-I model, Mono-I model, and Bi-I model, respectively. Arrows represent the vector field of the derivative of the first order moment and the orange line is the mean expression time course from an initial expression value (mRNA A, mRNA B)=(0, 0) to the time horizon *T* = 60. **d.** to **f.** show the mean, covariance and the skewness time courses, respectively, of the three interaction models. **g.** to **i.** show the pseudo-time reordering of the data corresponding to **a.** to **c.**, with the red line representing the pseudo-time mean.

To understand the origin of the crescent shape, we compared the time course of the skewness of mRNA A and mRNA B in the three GRN models. We found that in both the Bi-I and No-I model, the mRNA A was positively skewed and followed the identical time course. Furthermore, in the Bi-I model, mRNA B was also skewed similarly to mRNA A. That is, all downregulated mRNAs in the models exhibited similar skewness (Fig. 3 f, Sup. Fig. A c-d).

In summary, comparing only the mean time course of the three GRN models, we could not distinguish the underlying regulatory reactions between the No-I and Bi-I model. Similarly, the covariance could distinguish that Mono-I and Bi-I had some ‘negative’ interaction occurring, relative to the No-I model. However, the direction of the interactions was unclear. When we compared the skewness of the mRNAs, we could see that in the Mono-I interaction, mRNA A was being affected, and a similar effect was also acting on both mRNA A and B in the Bi-I model. Hence, the regulatory information was not in one statistic, but rather distributed over at least three statistics: the mean, the covariance, and the skewness.

#### 2.2.2 Pseudo-time augmented snapshots do not recapitulate the skewness in the original data

When multiple snapshots are unavailable, pseudo-time based augmentations of the data are used to infer the underlying GRNs. To study if the pseudo-time augmentation preserves the moments, we removed all the true time labels within each of the three GRN model’s data, and augmented the expression counts with a diffusion map based pseudo-time (see Section 4.5). In the time course of the central moments of the pseudo-time augmented data, we observed that both the No-I and Bi-I model’s mean expression of the mRNAs had a similar trend, like in the true time course (Fig. 3 g-i, Sup. Fig. A e). With respect to the covariance, we found that pseudo-time augmented data had negative covariance in the Mono-I and Bi-I models. Surprisingly, we found a positive covariance in the No-I model time augmented data (Sup. Fig. A f-g). The most drastic differences were seen in the skewness, where for all three models, the pseudo-time augmented data showed predominantly negative skewness, sharply contrasting against positive skewness seen in the original data. In summary, pseudo-time augmented data can capture trends in the first two central moments, however, it could underestimate the skewness, hindering accurate GRN inference.

#### 2.2.3 No- and Mono-directional interactions are harder to infer than Bi-directional interactions

Four inference methods were applied to infer GRNs from the synthetic scRNA-seq data of the three interaction models: the Mutual Information (MI) method (see Methods 4.6.1), the SINCERITIES method (see Methods 4.6.2), the linear moment based method (Linear MBI, see Methods 4.6.3), and the nonlinear moment based method (Nonlinear MBI, see Methods 4.6.4). We repeated the inference 400 times, each time generating from new synthetic data.

The MI method inferred non zero MI scores for all three models (see Method 4.6.1). We observed a more than five fold increase in the mean edge score for the Bi-I model with respect to the No-I model, and furthermore, the mean edge score for the Mono-I model was found in between (Supp. Fig. C a). A one-way ANOVA analysis showed that the differences in the means of the edge scores of the three models were statistically significant (an F-value of 49418 and a p-value of strictly less than 0.001). Furthermore, a pairwise comparison with Tukey HSD (with a p-value of 0.001) also showed a significant difference between each two models. Using the mean MI-score of the No-I model as the minimum score edge cutoff, we concluded that the MI based approach was effective in detecting that the three models had a different magnitude of interactions between the genes (Fig. 4 a).

**Figure 4:**
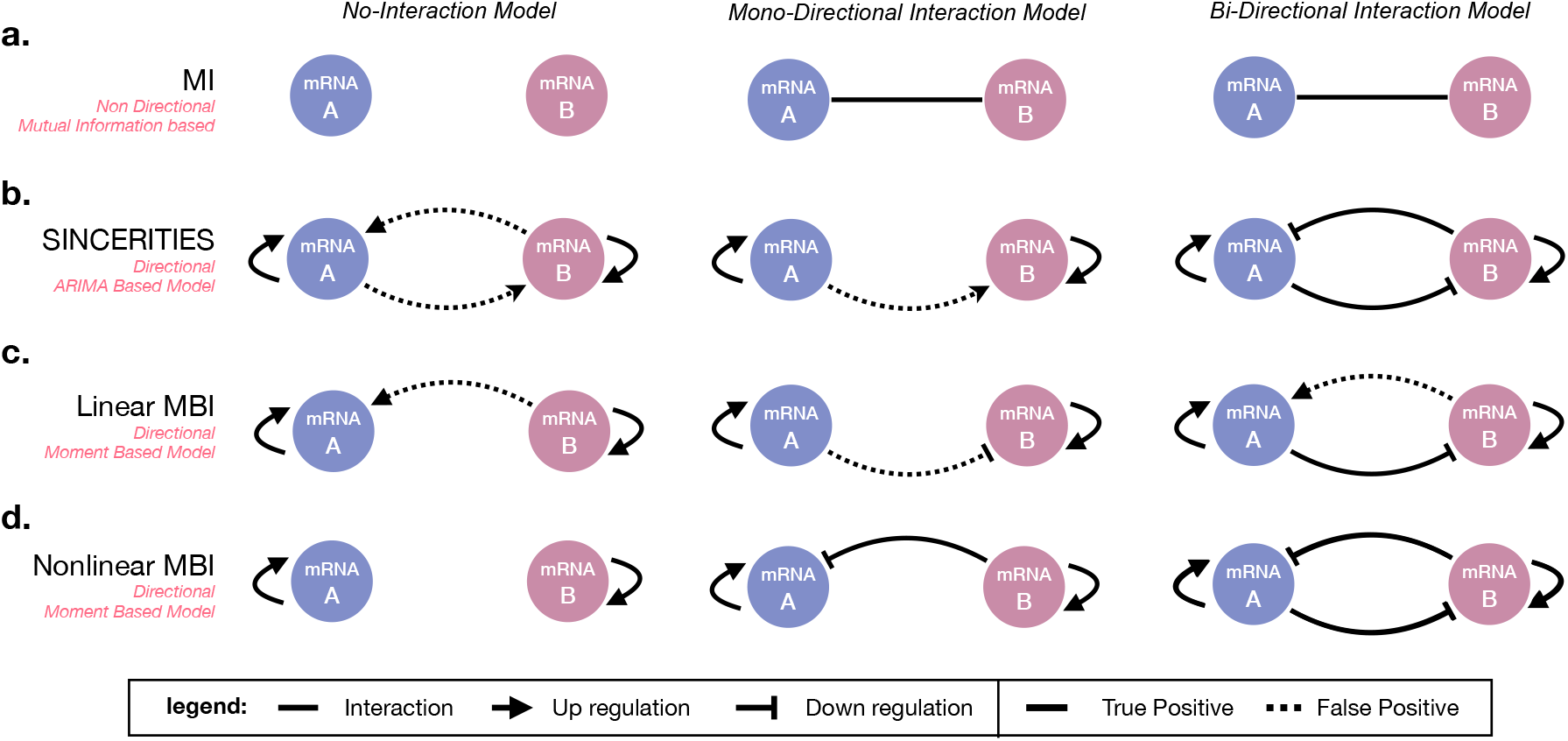
GRNs obtained from four inference methods. The inferred networks corresponding to the data from the three models No-I, Mono-I, and Bi-I, are columned left to right, respectively. The rows correspond to the inference method used: **a.** MI method, **b.** SINCERTIES, **c.** Linear MBI, and **d.** Nonlinear MBI.

In SINCERITIES, the interaction strength score is estimated by regularised regression of a system of distributional distances, while the sign of interaction (activation vs. repression) is determined by the sign of the partial correlation coefficient (see Method 4.6.1). In the No-I data, the SINCERITIES method inferred all possible activations between and within genes with a weak consistency in interaction scores (Fig. 4 b Left, Supp. Fig. C b). The interaction scores for the Mono-I model gave a clearer result, where a true positive self-activation of mRNA A and mRNA B were observed, however, the repression of mRNA A by mRNA B was missing. Instead, SINCERITIES inferred a false positive interaction of mRNA A activated by mRNA B (Fig. 4 b Middle). The inference of the Bi-I model was done correctly by SINCERITIES with high interaction scores (Fig. 4 b Right, Supp. Fig. C c).

The moment based inference (MBI) methods used the time course of up to degree three moments to infer their GRNs (see Methods 4.6.3–4.6.4). Given we knew *a priori* that the information was in the first three moments, to avoid over fitting, we used 15 times fewer snapshots in the moment based inference methods than in MI and SINCERITIES.

We saw that the Linear MBI performed slightly worse than SINCERITIES in inferring the underlying GRNs of the three models (Fig. 4 c). In particular, the Linear MBI method predicted a false positive activation between mRNA B and mRNA A in the Bi-I Model. Looking at individual GRNs inferred in the 400 replicates, we found that the Linear MBI method at best inferred the correct GRN 2.5 % of the time (Supp. Fig. B b).

Lastly, the Nonlinear MBI method performed the best out of the four methods. It predicted all true positive interactions and no false positive interactions (Fig. 4 d). Furthermore, looking to the individual GRNs in the replicates, we found that it correctly predicted the No-I model 54 % of the time, the Mono-I model 78 % of the time, and the Bi-I model 95 % of the time (Supp. Fig. B c).

In summary, in the four inference methods that we compared, the Bi-I model was the easiest to capture (Fig. 3 c). We suspect that this results from the strong double correlation signal present in the data, which results from the nonlinear interaction between Gene A and Gene B. The fact that the Mono-I model only had one interaction was detected by all methods, however, the directionality and regulatory mechanism could not be correctly detected. Lastly, the No-I model showed that not all methods are specific enough to correctly detect no interaction.

### 2.3 Linear MBI methods are sensitive to interval lengths between snapshots

The linear least-squares method is well established and can be used to solve high-dimensional inference problems. To harness its scalability for inferring GRNs using moments (Linear MBI), good approximations of the derivatives of the moments’ time courses are essential (see cartoon in Fig. 5 a). However, due to snapshot intervals generally being large in sequencing experiments, good derivative approximations are seldom possible. We investigated the effect of interval lengths between snapshots on the accuracy of the inference by constructing a simple stochastic damped oscillator model (see Methods 4.8, Fig. 5 b-c). Snapshots of different time interval lengths were taken and their underlying network was inferred using the Linear MBI method and the Nonlinear MBI method.

**Figure 5:**
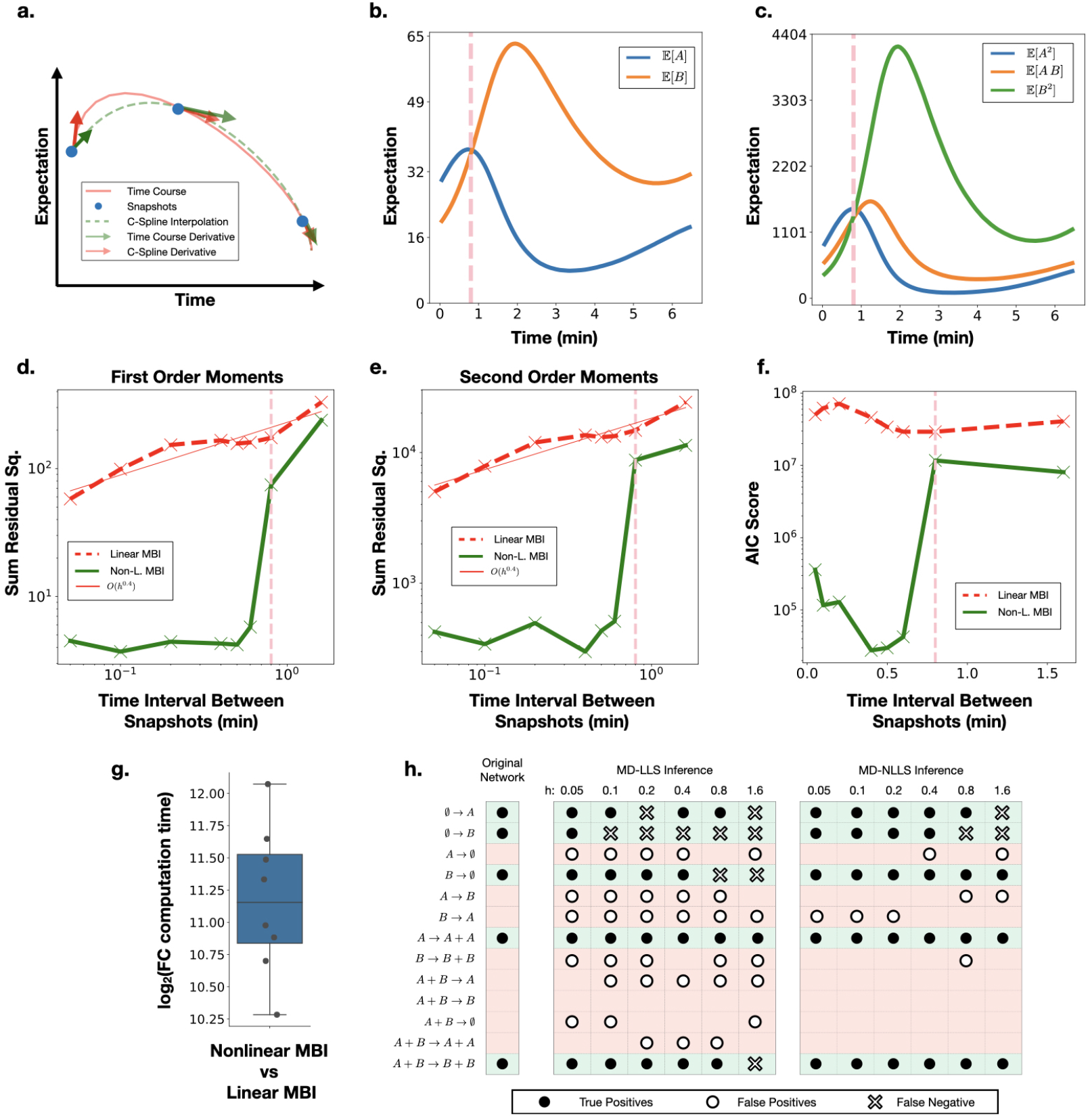
Performance of MBI methods. The performance of Linear MBI and Nonlinear MBI methods on the stochastic damped oscillator model. **a.** Cartoon showing the interpolation and derivative estimation of snapshot time courses. **b-c.** Time course of the first and the second order moments, respectively. **d-e.** Residual sum of squares of the first and second order moments for varying snapshot interval lengths, respectively. **f.** AIC score for varying snapshot interval lengths. The linear MBI (red dashed line), the non-linear MBI (green solid line) are shown. **b-f.** The first peak in the mean time course of population A (pink dashed vertical line). **d-e.** The linear fit of the residual least squares of the linear MBI method (thin red line). **g.** Log 2 fold change in computation time non-linear MBI vs linear MBI. **h.** The true positive (solid dot), false positive (hollow dot), and the false negative (hollow cross) reactions inferred by the linear and the non-linear MBI methods for varying snapshot interval lengths of the stochastic oscillator model.

### 2.4 The Linear MBI method struggles even at small snapshot intervals

We observed that the residual sum of squares of the Linear MBI method increased with order 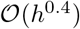 with respect to interval length *h* between snapshots (Fig. 5 d-e). Upon inspecting the inferred reactions, we found that for time interval of *h* = 0.05 min, the Linear MBI method inferred the five true reactions and a further five false positive reactions (Fig. 5 h). For the subsequent interval lengths, we found that the Linear MBI method continued inferring five to six false positive reactions and the number of true positive reactions was decreasing.

The mean time course of the SSA simulations with the inferred parameters showed that the Linear MBI method performed poorly in fitting data, even for the smallest interval length of *h* = 0.05 min (Supp. Fig. E). In this case, 128 snapshots (1152 moments) were used to infer 13 reactions and surprisingly, we did not observe a close reconstruction of the real data. This suggests that the errors made in estimating the derivative could not be remedied by the large amount of snapshot data.

#### 2.4.1 The nonlinear MBI method circumvents the derivative estimation step at the cost of a significant increase in computational time

The Nonlinear MBI method circumvents the derivative estimation by minimising the distance of the inferred model to the data. This results in a non-linear least squares problem, which does not need the time-course derivative of the moments. In comparison to the Linear MBI method, for time intervals less than *h* = 0.6 min, we found that the Nonlinear MBI method had at least one order of magnitude lower residual sum of squares in all moments (Fig. 5 d-e). Furthermore, we found that the residual sum of squares did not increase linearly for small time interval lengths, showing a near flat trend between residual and interval length. Looking at the inferred reaction network, we observed that the Nonlinear MBI method captured all of the true reactions, and only inferred one false positive reaction for time intervals less than *h* = 0.6 min (Fig. 5 h). Interestingly, we observed that for intervals larger than *h* = 0.8 min, the Nonlinear MBI method starts to perform as poorly as the Linear MBI method. Upon closer inspection, we found that *h* = 0.8 min is roughly where the first peak in the time course of population A occurs (Fig. 5 b-c). Comparing the AIC scores of the two approaches, we saw that the Nonlinear MBI’s minimum AIC score was at least two orders of magnitude smaller than that of the Linear MBI (Fig. 5 f). Even though the Nonlinear MBI method performed better, it must be noted that it took on average nearly 2000 times longer to compute than the Linear MBI method (Fig. 5 g).

In summary, the simple stochastic damped oscillator model highlighted the major challenges of using moments based methods for inference. In particular, we observed that the log of the residual scaled sublinearly with the interval length for the Linear MBI method. In order to achieve a similar accuracy as the Nonlinear MBI method at interval length *h* = 0.05 min, the interval length of the data for the Linear MBI would have to be smaller than 10^*−*3^ min. Furthermore, we saw that the snapshot interval has to be small enough to observe the turning points of the system for accurate inference.

## 3 Discussion

We considered three GRN models which contained a varying mixture of correlation and regulation between the mRNA populations. The highest degree of correlation was contained in the No-interaction model, a medium mixture of correlation and regulation was contained in the Mono-interaction model, and the highest degree of regulation was contained in the Bi-directional model. We can draw three key conclusions from the experiments conducted in this paper. The first being that the at least up to the third-order moments are required for GRN inference. The second being that only the nonlinear moment-based inference method is able to consistently infer the ground truth GRN of our artificial scRNA-seq data, thereby a high potential to infer GRN of real-world data. And the third being that certain data pre-processing steps like pseudotime are not recommended. These three key conclusions are discussed in more details bellow.

Our experiments allowed us to conclude that at least up to the third-order moments are essential for GRN inference. We saw that regulatory reactions in our models generated distinctive signatures in the higher-order (second- and third-order) moments of the dataset. These signatures, however, could not fully be detected by the distribution-based inference methods (MI and SINCERITIES), because the summary statistics that they utilised were not related to the higher-order moments, which led to an under-performance of these inference methods. The MI method is based on threshold strategies which could detect interacting genes, however, it could not infer the direction of the interaction, and smaller interactions are likely to be false negatives due to the threshold cut-off. Similarly, the SINCERITIES method is also based on threshold strategies with the difference that it infers the direction of the regulation. However, the summary statistics used by SINCERITIES falsely detected the correlation as regulation in the No-I and Mono-I models, suggesting that it was not able to distinguish the nuances between correlation and regulation. The Linear moment-based inference method performed poorly despite using the moments’ derivatives as a summary statistic, we speculate that its under-performance is a consequence of the time-interval between the data snapshots not being ideal for the derivative estimation. In contrast to the other methods, the nonlinear moment-based inference method was able to consistently reconstruct our three GRN models.

Using higher-order moments has three key impacts on the inference of GRNs from scRNA-seq data: firstly, higher-order moments can clearly distinguish between correlation and regulation, which is due to regulatory information being present across the higher-order joint moments, unlike correlation. As a consequence, fewer false positive and -negative interactions are inferred, leading to more true positive regulatory reactions. Secondly, a summary statistic that captures regulation should be based on the moments, for example, if it satisfies Taylor’s theorem, then it can be arbitrarily-well approximated by a polynomial of the moments. Lastly, to use higher-order moments, the synthetic data model has to be redesigned to be a Markov jump process, incorporating Poisson intrinsic noise and moving away from mean-driven Gaussian noise.

The higher-order moments proved to be robust summary statistics for regulatory interactions, however, they might not be conserved during data pre-processing. Additionally, the fact that they evolve through time non-linearly causes a major computational challenge for the inference. Thus, new numerical schemes to solve high dimensional non-linear least squares problems are essential in furthering the field of inferring GRNs.

In regards to limitations, our synthetic models were designed to be ideal, with respect to extrinsic noise, to strongly focus on detecting the regulatory signatures and to study how to extract this signature. However, in practice, real biological data have far more caveats which need to be captured by the synthetic data. Similarly to the synthetic data generator SERGIO (36), we envisage to adapt our Markov-jump model to simulate more realistic synthetic data, including technical issues such as drop-outs, Hill’s function propensities, multi-gene interactions, and cell cycles.

A major obstacle for finding good summary statistics (feature extraction) has been the lack of ground truth GRNs for scRNA-seq datasets (11; 37). True GRNs that were generated from bulk RNA-seq experiments have not been reproduced with single-cell experiments (38). It is current practise to use protein interaction networks to check for false positive interaction in the GRNs inferred from scRNA-seq data (39). Since protein interaction networks only indicate whether a gene is being expressed or several genes are being co-expressed, they are good at filtering genes which are correlated, however, the underlying regulation and directionality is not captured. Additionally, ChiP-seq datasets provide potential biding sites of transcription factors, however, given a transcription factor has multiple binding sites and can also form complexes, we can establish the directionality of regulation, but lose specificity. In conclusion, we advocate that ground truth datasets require multi-omics single cell datasets (37). Such datasets are imperative for calibrating current- and designing new GRN inference methods based on single cell technology.

## 4 Methods

### 4.1 Trends in the mean and variance of gene expression datasets

We considered three different single cell snapshot datasets generated from laboratory experiments. These datasets were: (1) a scRNA-seq dataset from human embryonic stem cells (hESC) from which we removed genes with more than 50% drop out cells to get a dataset of 8917 genes and 6 snapshot measurements (40), (2) a qRT-PCR dataset from THP-1 cells containing 48 genes and 8 snapshots (41), and (3) a qRT-PCR dataset from mouse embryonic stem cells (mESC) containing 500 genes and 7 snapshots (42).

We also considered two synthetic datasets used for benchmarking GRN inference methods. One dataset was generated by GeneNetWeaver (GNW) (43; 44). It contained 1000 time series with 21 snapshots of *E. coli* gene expression of 500 genes. The other dataset was obtained using the BoolODE model (11). We selected the simulation of “linear long” GRN topology which consisted of 18 genes. This dataset included ten replicate simulations, each containing gene expression measurements for 5000 cells and the time at which they were sampled. We pooled the replicate simulations to obtain 1499 snapshots of the gene expression of 50000 cells.

For each dataset, the time course of the mean and variance of gene expression was computed. We then normalised the time courses of the mean and variance of gene expression to values in the interval (−1, 1) by shifting each time point value by the first value in the time course, and then dividing the shifted values by the maximum magnitude of gene expression of each gene. We further performed PCA on the normalised time courses and projected the datasets onto the fewest number of principal components explaining around 95% of the total variation.

The projected datasets were then clustered into trends using the k-means clustering algorithm in which the number of clusters was chosen according to a discrete uniform distribution between 4 and 8. Finally, we estimated the marginal and joint probability distribution of a gene being associated to a trend in the mean and being associated to a trend in the variance. These probabilities were used to compute the proficiency measure representing the proportion of trends in the variance that can be explained by the trends in the mean. We repeated this procedure 1000 times to obtain a proficiency distribution for each of the datasets.

### 4.2 Two Gene Interaction Model

The interaction networks in the three models No-I, Mono-I, Bi-I, were designed to have a nested property, that is, the No-I model was a sub-network of the Mono-I model and the Mono-I model was a sub-network of the Bi-I model. At any time, a gene has a binary state space {**on**, **off**}. The mRNA and proteins are described by their counts, therefore they have a positive integer state space. The interaction model considered two genes: Gene A and Gene B, and the species that were involved in the reactions are Gene A, Gene B, Protein A, Protein B, mRNA A, and mRNA B.

Table 1 describes each component of the No-I model. The Mono-I interaction model contains all reactions in Table 1 with the exception that Reaction 1 is replaced with Reaction 1a (Table 2) to involve one of the protein products (here Protein B) which actively up-regulates the switching off of Gene A, and Reaction 5 is modified to Reaction 5a. The Bi-I interaction model also contains all reactions in Table 1, with the changes shown in Table 3, i.e., Reaction 1 is replaced with Reaction 1a as in the Mono-I interaction model, and Reaction 2 is replaced with Reaction 2a to include the up-regulation of the switching off of Gene B by Gene A, and the propensity coefficients of Reactions 5 and 6 are modified to that of Reaction 5a and 6a.

**Table 1:** Components of the two-gene No-Interaction model. The positions in the stoichiometry vector correspond to (Gene A, Gene B, Protein A, Protein B, mRNA A, mRNA B)

| # | Reactions | Coeff. ($\text{min}^{-1}$) | Propensities | Stoichiometry | Description |
| --- | --- | --- | --- | --- | --- |
| 1 | Gene A **on** ⟶ Gene A **off** | $\sigma_1 = 0.125$ | $\sigma_1 [\text{Gene A on}]$ | $(-1, 0, 0, 0, 0, 0)$ | Inactivation |
| 2 | Gene B **on** ⟶ Gene B **off** | $\sigma_2 = 0.125$ | $\sigma_2 [\text{Gene B on}]$ | $(0, -1, 0, 0, 0, 0)$ | Inactivation |
| 3 | Gene A **off** ⟶ Gene A **on** | $\sigma_3 = 0.5$ | $\sigma_3 [\text{Gene A off}]$ | $(1, 0, 0, 0, 0, 0)$ | Activation |
| 4 | Gene B **off** ⟶ Gene B **on** | $\sigma_4 = 0.5$ | $\sigma_4 [\text{Gene B off}]$ | $(0, 1, 0, 0, 0, 0)$ | Activation |
| 5 | Gene A **on** ⟶ Gene A **on** + mRNA A | $\rho_1 = 4.75$ | $\rho_1 [\text{Gene A on}]$ | $(0, 0, 0, 0, 1, 0)$ | Transcription |
| 6 | Gene B **on** ⟶ Gene B **on** + mRNA B | $\rho_2 = 4.75$ | $\rho_2 [\text{Gene B on}]$ | $(0, 0, 0, 0, 0, 1)$ | Transcription |
| 7 | Gene A **off** ⟶ Gene A **off** + mRNA A | $\rho_3 = 1.0$ | $\rho_3 [\text{Gene A off}]$ | $(0, 0, 0, 0, 1, 0)$ | Transcription |
| 8 | Gene B **off** ⟶ Gene B **off** + mRNA B | $\rho_4 = 1.0$ | $\rho_4 [\text{Gene B off}]$ | $(0, 0, 0, 0, 0, 1)$ | Transcription |
| 9 | mRNA A ⟶ mRNA A + Protein A | $\gamma_1 = 5.0$ | $\gamma_1 [\text{mRNA A}]$ | $(0, 0, 1, 0, 0, 0)$ | Translation |
| 10 | mRNA B ⟶ mRNA B + Protein B | $\gamma_2 = 5.0$ | $\gamma_2 [\text{mRNA B}]$ | $(0, 0, 0, 1, 0, 0)$ | Translation |
| 11 | Protein A $\longrightarrow \emptyset$ | $\kappa_1 = 0.1$ | $\kappa_1 [\text{Protein A}]$ | $(0, 0, -1, 0, 0, 0)$ | Degradation |
| 12 | Protein B $\longrightarrow \emptyset$ | $\kappa_2 = 0.1$ | $\kappa_2 [\text{Protein B}]$ | $(0, 0, 0, -1, 0, 0)$ | Degradation |
| 13 | mRNA A $\longrightarrow \emptyset$ | $\delta_1 = 0.1$ | $\delta_1 [\text{mRNA A}]$ | $(0, 0, 0, 0, -1, 0)$ | Degradation |
| 14 | mRNA B $\longrightarrow \emptyset$ | $\delta_2 = 0.1$ | $\delta_2 [\text{mRNA B}]$ | $(0, 0, 0, 0, 0, -1)$ | Degradation |

**Table 2:** Reactions to be replaced in Table 1 to obtain the Mono-Interaction model. The positions in the stoichiometry vector corresponds to (Gene A, Gene B, Prot. A = Protein A, Prot. B = Protein B, mRNA A, mRNA B)

| # | Reactions | Coeff. ($\text{min}^{-1}$) | Propensities | Stoichiometry | Description |
| --- | --- | --- | --- | --- | --- |
| 1a | Gene A **on** + Prot. B ⟶ Gene A **off** | $\sigma_1 = 0.01875$ | $\sigma_1 [\text{Prot. B}][\text{Gene A on}]$ | $(-1, 0, 0, -1, 0, 0)$ | Inactivation |
| 5a | Gene A **on** ⟶ Gene A **on** + mRNA A | $\rho_1 = 6.0$ | $\rho_1 [\text{Gene A on}]$ | $(0, 0, 0, 0, 1, 0)$ | Transcription |

**Table 3:** Reactions to be replaced in Table 1 to obtain the Bi-Interaction model. The positions in the stoichiometry vector correspond to (Gene A, Gene B, Prot. A = Protein A, Prot. B = Protein B, mRNA A, mRNA B)

| # | Reactions | Coeff. ($\text{min}^{-1}$) | Propensities | Stoichiometry | Description |
| --- | --- | --- | --- | --- | --- |
| 1a | Gene A **on** + Prot. B ⟶ Gene A **off** | $\sigma_1 = 0.01875$ | $\sigma_1 [\text{Prot. B}][\text{Gene A on}]$ | $(-1, 0, 0, -1, 0, 0)$ | Inactivation |
| 2a | Gene B **on** + Prot. A ⟶ Gene B **off** | $\sigma_2 = 0.01875$ | $\sigma_2 [\text{Prot. B}][\text{Gene B on}]$ | $(0, -1, -1, 0, 0, 0)$ | Inactivation |
| 5a | Gene A **on** ⟶ Gene A **on** + mRNA A | $\rho_1 = 6.0$ | $\rho_1 [\text{Gene A on}]$ | $(0, 0, 0, 0, 1, 0)$ | Transcription |
| 6a | Gene B **on** ⟶ Gene B **on** + mRNA B | $\rho_2 = 6.0$ | $\rho_2 [\text{Gene B on}]$ | $(0, 0, 0, 0, 0, 1)$ | Transcription |

### 4.3 Synthetic scRNA-seq data

For each of the two-gene interaction models No-I, Mono-I, and Bi-I, species counts of (Gene A, Gene B, Protein A, Protein B, mRNA A, mRNA B) were generated using the stochastic simulation algorithm (SSA) (45). The initial population configuration was set to (Gene A **on**, Gene B **on**, 0, 0, 0, 0, 0) to mimic the accessibility of the gene on the chromatin, and the initial simulation time was set to 0 min. Gene states and populations counts in the simulations were sampled in time intervals of 0.5 min and up to a time horizon of 60 min. A sample snapshot of SSA mRNA population counts can be interpreted as scRNA-seq data of an individual cell belonging to a fixed cell population. For each model, we generated a total of 100000 time trajectories of mRNA species counts, thus simulating temporal snapshots of scRNA-seq data of each individual cell for a population of 100000 cells. We then constructed 400 replicates of this mRNA count dataset. In each replicate, we subsampled without replacement 10000 time trajectories from the 100000 cells. The underlying GRNs schematics were inferred according to specific rules suited to each of the GRN inference methods that we applied (see Section 4.7).

### 4.4 Computing the moments

The moments of the snapshot mRNA population were required for moment-based GRN inference methods (Section 4.6.3 and 4.6.4). Let the population of mRNA A in the *d*-th snapshot of the *n*-th cell be denoted by *a*_*d*,*n*_, similarly for mRNA B, *b*_*d*,*n*_. Then for any pair of non-negative integers (*l*_1_*, l*_2_) such that *l* = *l*_1_ + *l*_2_, the *l*-th order moment of the *d*-th snapshot is given by,

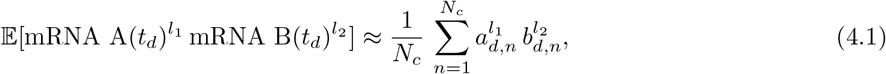

where *N*_*c*_ is the total number of cells in the snapshot sample.

### 4.5 Pseudo-time ordering of the data

We performed pseudo-time ordering using diffusion maps as described in (29) and implemented in SCANPY (46). This technique is used to achieve a temporal ordering of non-ordered snapshot observations of RNA-seq gene expression (8; 9; 27; 47). The diffusion map yielded a non-linear dimension reduction and a denoised representation of the high dimensional gene expression data. The pseudo-time of a cell was defined as the measured distance from a given root cell which was assigned a pseudo-time of zero a priori. The pseudo-time ordered data was suitable for lineage branching inference due to the ability of diffusion maps to recover the mean dynamics of gene expression (29). Furthermore, the application of the pseudo-time method has been extended to infer GRNs (27; 48).

We removed all true time labels in the synthetic mRNA count data (Section 4.3) and created a typical pseudo-time ordering on subsampled snapshots as follows: We subsampled from the 100000 simulated trajectories representing temporal snapshots of mRNA counts for 100000 cells. We slightly oversampled cells from earlier time points, as our simulation of gene expression converges to equilibrium in later time points, which would bias the pseudo-time approach. Furthermore, we excluded the first snapshot at time 0 min. We generated three datasets comprising the expression of two genes for 50000 cells. An arbitrary cell from the first time point was chosen to be the root cell of the diffusion pseudo-time ordering. The pipeline was implemented in SCANPY by computing a sparse nearest neighbour graph (50 neighbours), the observations were embedded in diffusion map space of three dimensions, and the diffusion pseudo-time for each cell was computed using the first two dimensions.

### 4.6 GRNs inference methods

We inferred GRNs from the synthetic scRNA-seq data (Section 4.3) representing temporal snapshots of mRNA counts of each of the two-gene interaction models described in Section 4.2. Since the mRNA counts are the only information available, we are not aiming to reconstruct the two-gene interaction models. Instead, we aim to capture the regulatory relationships between the genes reflected through the interaction of the mRNAs.

#### 4.6.1 MI method

The mutual information (MI) measure quantifies the amount of information shared between two discrete random variables *X* and *Y* and is formulated as follows,

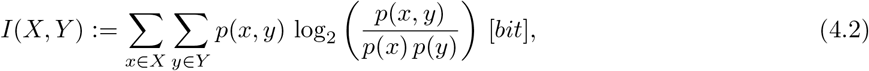

where *p*(*x*) and *p*(*y*) are the probability distributions of *X* and *Y* , respectively; and *p*(*x, y*) is the joint probability distribution of *X* and *Y* .

The MI is a symmetric measure, therefore it has been used to infer non-directed GRN by using it as a score for the confidence of an edge between the genes (22). Since MI can take any positive value, there is no general way of interpreting it’s magnitude. A threshold of top scoring network is usually used to infer the underlying networks.

The mRNA count datasets, generated from our two-gene interaction model, were used in the Mutual Information (MI) GRNs inference method. The MI scores were then computed using the implementation provided by Chan et al (22). Additionally, it was not possible to choose a number of top scoring networks because there was only one possible edge given by the MI method. Therefore, we developed a different way of interpreting inferring the underlying GRNs (see Section 4.7.1).

#### 4.6.2 SINCERITIES method

SINCERITIES (SINgle CEll Regularized Inference using TIme-stamped Expression profileS) has recently been proposed to infer directed GRNs by using temporal snapshots of gene expression data (24). The SINCERITIES algorithm implements a regularised regression of a system of Kolmogorov–Smirnov distributional distances and assigns a ranked list of scores 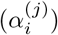 representing the influence of gene *j* on any other gene *i* in the dataset. A large score indicates a higher confidence that the corresponding edges exists. Furthermore, SINCERITIES infers the direction of the edges, that is, the nature of the interaction, from the sign of the partial correlation coefficients between each two genes. Similarly to the MI method (see Section 4.6.1), there exists no general way of interpreting the magnitude of the 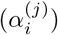 scores. As the authors of SINCERITIES leave it to the user to choose a score threshold for drawing an edge between the genes, we chose a generous score cutoff and inferred the underlying GRNs according to specific rules (see Section 4.7.1).

#### 4.6.3 Linear MBI method

In the Linear Moment Based Inference (MBI) method, we used an adaptation of the SINDy method (Supp. Sec. A.2) to infer GRNs from the data (26). We modelled GRNs as Markov jump processes (see Supp A.1), representing the time evolution of *N_s_* species undergoing *N_r_* chemical reactions. Therefore, we aimed to discover the linear dynamical system that defines the moments of the mRNA species counts involved in the reaction network (Supp. Eq. A.4).

We thus considered the state vector 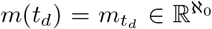 containing the *raw moments* of the *N_s_* species at time *t_d_* for *d* ∈ {1*, … . D*}, and 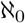 denotes the cardinality of the countably infinite set ℕ. Then, we constructed a library of candidate non-linear functions Ψ(**M**) as follows,

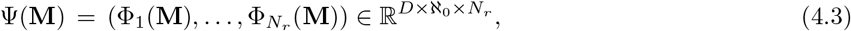

where 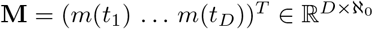, and

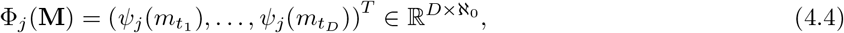

is the vector of *stoichiometric moment functions* of reaction *i* (26) which is given as follows,

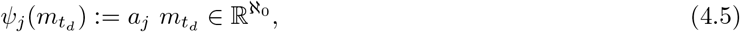

where *a*_*j*_ is the *design block* corresponding to reaction *i* in the *design matrix* **A** of the moments equations (Supp. Eq. A.4). This allowed us to formulate the following linear system,

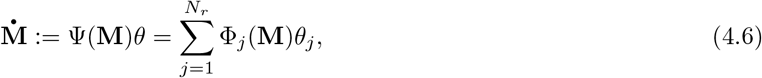

where 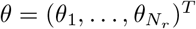 is the vector of propensity coefficients of the *N*_*r*_ reactions that we considered.

A finite dimensional approximation of Eq. 4.6 was obtained by truncation of the moments vector. According to Supp. Eq. A.3, up to order *l* +1 moments are required to model the time derivatives of up to order *l* moments. Thus, we can formulate the truncated system,

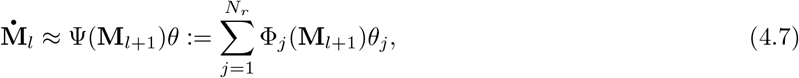

where **M**_*l*_ = (*m*_*l*_ (*t*_1_) … *m*_*l*_ (*t*_*D*_))^*T*^ ∈ ℝ*D×l* with *m*_*l*_ being the vector of up to order *l* moments.

For the GRN inference, we aimed to discover a sparsely connected network which reflects the minimal set of reactions that are involved in the network (26; 49), because it has been demonstrated that robust GRNs are parsimonious (50). Therefore, sparse regression minimisation techniques are applied to find the sparse parameter vector 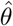 satisfying

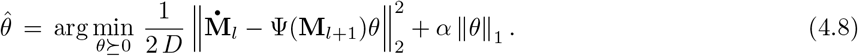

This problem was approximated by

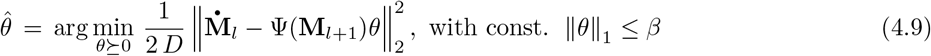

where *β* is the upper bound on the sum of the parameters and was set to 1000 in our calculations. In our simulations, we used up to order four moments of the synthetic mRNA counts, i.e., solving Eq. 4.8 for *l* = 3. In addition, we removed the first 30 data points for a more accurate representation of the moments and further reduced the dataset by 15 times to avoid overfitting. The problem in Eq. 4.9 was solved as a quadratic program (51). Solutions were obtained using python. The underlying GRNs were then generated according to specific rules (see Section 4.7.2).

#### 4.6.4 Non-linear MBI method

Similarly to the Linear Moment Based Inference (MBI) method, the Non-linear MBI method also inferred the parameters of the reaction kinetics of a Markov jump process from the moments equation (Supp. Eq. A.1). Since the moments equation is an infinite-dimensional system, we handled the truncation problem by introducing the interpolations of the higher-order moments from the data source into a truncated system.

We considered *m*_*l*_ as the vector containing moments up to order *l*. We then approximated the moments equation with,

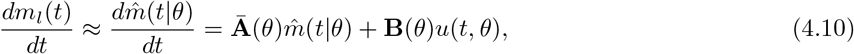

where 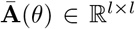 is linear in the parameter *θ*, and is the block of the design matrix **A** that comprises the dependency of the derivatives of order *l* moments on themselves (Supp. Eq. A.4). **B** is a rectangular matrix of dimensions (dim 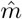, dim *u*(*t*)), and is the block of the design matrix **A** that includes the dependency of the derivatives of order *l* moments on the order *l* + 1 moments, and *u*(*t*, *θ*) is the vector interpolation of the moments of order *l* + 1 from the data.

Given *D* temporal snapshots of moments data 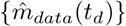 with *d* ∈ {1, … . *D*}, the parameter 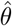 that best represents the data in the model Eq. 4.10 can be obtained via maximum likelihood estimation. If we assume that the errors in the moments model are normally distributed, 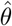 can be obtained by minimising the negative log-likelihood function which is proportional to,

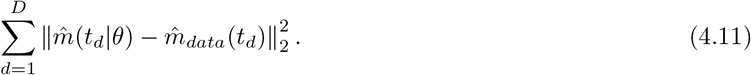

The problem is then reduced to the non-linear least-squares minimisation problem,

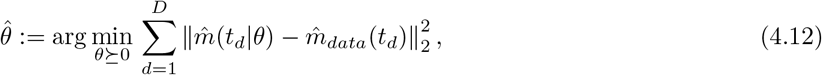

where 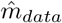 is the vector containing moments up to order *l* of the data.

We implemented the Non-linear MBI method in python using SciPy (52). We used the same moments datasets as in the Linear MBI method (see Section 4.6.3), which included up to the fourth order moments of the synthetic mRNA counts data (see Section 4.3). The vector 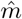 contained up to order *l* = 2 moments. We computed the splines *u* of the order *l* + 1 = 3 moments by specifying the moments’ derivative (Eq. 4.10) at the end points of the temporal snapshot data, i.e., at *t*_0_ and *t*_*D*_ (this required the moments of order *l* + 2 = 4).

A numerical approximation of 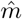 was generated by solving Eq. 4.10 for each two adjacent time point snapshots and fixed *θ*, that is, by solving the initial value problem,

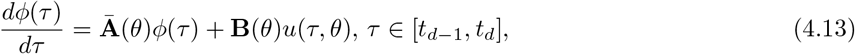

with the initial condition 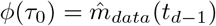 for each *d* ∈ {2*, … . D*}.

The solution of Eq. 4.13 yielded the approximation 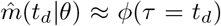. This approximated solution was then used to compute the error function in Eq. 4.11. Due to a large difference between the magnitude of order one (the means) and order two moments, we multiplied the error in the mean by a constant weight of 40 to avoid its underestimation. A standard least squares minimisation routine was used to find 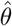 solving Eq. 4.12. The underlying GRN was then inferred using a set of rules that we defined in Section 4.7.2.

### 4.7 GRN Schematics

#### 4.7.1 Score-based schematics: MI and SINCERITIES methods

We conducted an ANOVA analysis of the MI score distribution of the three two-gene interaction models No-I, Mono-I, and Bi-I. We then used the mean MI score of the No-I model as a minimum score edge cutoff for the inferred GRNs.

For the SINCERITIES method (Section 4.6.2), the algorithm generated scores that represent the confidence of an edge between two genes. We then chose a generously low threshold of 0.05 to draw an edge. Additionally, the SINCERITIES method provided the direction of the edge. Therefore, we were able to infer GRNs directly from the output of SINCERITIES. We used the algorithm to infer a GRN for each of the 400 replicates of the mRNA count dataset (Section 4.3). The frequencies of the different resulting interaction networks across the 400 runs are depicted in Supp. Fig. 2 a, and we chose the most frequently inferred network as the underlying GRN (Fig. 3).

#### 4.7.2 Flux-based schematics: MBI methods

We set up the Linear and Non-linear MBI methods to infer the reaction network depicted in Table 4. Reactions 6 and 8 represent the up-regulation of A by B and B by A, respectively, while reactions 9 and 10 represent the down-regulation of A by B and vice versa. We then generated the following adjacency matrix,

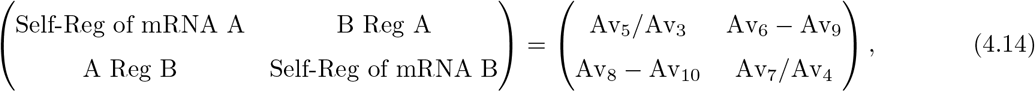

where Av_*j*_ are the average number of times reaction *j* fired within the time window. The diagonal of the adjacency matrix (Eq. 4.14) represents the average number of self-catalysis vs. degradation for mRNA A and mRNA B. We only considered a non-zero Self-Reg value if the average self-catalysis exceeded 10 within the time window. Since we only aimed to identify self-regulation, the ratio between the self-catalysis and the degradation reaction is used to obtain a positive number, but subtraction could be used instead of a ratio if we wanted to distinguish between self-upregulation (positive values) or self-downregulation (negative values).

**Table 4:** Reaction Library for two mRNA species interaction. For simplicity we refer to mRNA A as A and to mRNA B as B.

| # | Reactions | Propensity | Stoichiometry | Description |
| --- | --- | --- | --- | --- |
| 1 | $\emptyset \longrightarrow A$ | $\theta_1$ | $(1, 0)$ | synthesis of A |
| 2 | $\emptyset \longrightarrow B$ | $\theta_2$ | $(0, 1)$ | synthesis of B |
| 3 | $A \longrightarrow \emptyset$ | $\theta_3 A$ | $(-1, 0)$ | degradation of A |
| 4 | $B \longrightarrow \emptyset$ | $\theta_4 B$ | $(0, -1)$ | degradation of B |
| 5 | $A \longrightarrow A + A$ | $\theta_5 A$ | $(1, 0)$ | self catalysis of A |
| 6 | $B \longrightarrow A + B$ | $\theta_6 B$ | $(1, 0)$ | synthesis of A by B |
| 7 | $B \longrightarrow B + B$ | $\theta_7 B$ | $(0, 1)$ | self catalysis of B |
| 8 | $A \longrightarrow A + B$ | $\theta_8 A$ | $(0, 1)$ | synthesis of B by A |
| 9 | $A + B \longrightarrow B$ | $\theta_9 AB$ | $(-1, 0)$ | annihilation of A from encounter with B |
| 10 | $A + B \longrightarrow A$ | $\theta_{10} AB$ | $(0, -1)$ | annihilation of B from encounter with A |

In Eq. 4.14, A Reg B represents up-regulation vs. down-regulation for mRNA B by mRNA A, and vice versa for B Reg A. Thus, the sign of these entries determines the direction of the regulation, that is, whether it is a net up- or down-regulation. We only considered a non-zero regulation value if the net regulation was higher than ten times within the time window.

### 4.8 Evaluation of MBI method accuracy

We investigated the accuracy of the MBI methods by using the Stochastic Damped Oscillator (SDO) model. The interactions involved in the SDO are pictured in Table 5. Using the stochastic simulation algorithm (SSA), we generated synthetic population counts starting from an initial population count of (0, 0). The simulations were performed from an initial time of 0 min to a final time of 6.45 min by taking sample population counts at every time intervals of 0.05 min. The moments data, up to order four, were computed from 10000 SSA trajectories, which yielded *D* = 130 snapshot data points.

**Table 5:** Stochastic Damped Oscillator Reaction Library

| # | Reactions | Coefficients | Stoichiometry | Description |
| --- | --- | --- | --- | --- |
| 1 | $\emptyset \longrightarrow A$ | $\theta_1 = 4.0$ | $(1, 0)$ | birth of A |
| 2 | $\emptyset \longrightarrow B$ | $\theta_2 = 3.0$ | $(0, 1)$ | birth of B |
| 3 | $A \longrightarrow \emptyset$ | $\theta_3 = 0.0$ | $(-1, 0)$ | death of A |
| 4 | $B \longrightarrow \emptyset$ | $\theta_4 = 0.7$ | $(0, -1)$ | death of B |
| 5 | $A \longrightarrow B$ | $\theta_5 = 0.0$ | $(-1, 1)$ | transition of A to B |
| 6 | $B \longrightarrow A$ | $\theta_6 = 0.0$ | $(1, -1)$ | transition of B to A |
| 7 | $A \longrightarrow A + A$ | $\theta_7 = 1.25$ | $(1, 0)$ | self production of A |
| 8 | $B \longrightarrow B + B$ | $\theta_8 = 0.0$ | $(0, 1)$ | self production of B |
| 9 | $A + B \longrightarrow A$ | $\theta_9 = 0.0$ | $(0, -1)$ | annihilation of B from encounter with A |
| 10 | $A + B \longrightarrow B$ | $\theta_{10} = 0.0$ | $(-1, 0)$ | annihilation of A from encounter with B |
| 11 | $A + B \longrightarrow \emptyset$ | $\theta_{11} = 0.0$ | $(-1, -1)$ | annihilation of A and B from encounter |
| 12 | $A + B \longrightarrow A + A$ | $\theta_{12} = 0.04$ | $(1, -1)$ | birth of A from encounter with B |
| 13 | $A + B \longrightarrow B + B$ | $\theta_{13} = 0.04$ | $(-1, 1)$ | birth of B from encounter with A |

#### 4.8.1 Sensitivity of the MBI methods to interval lengths between snapshot

We investigated the effect of the time interval separation ∆*t* ∈ {0.05, 0.1, 0.2, 0.4, 0.5, 0.6, 0.8, 0.9, 1.6} between snapshots by inferring the reaction network parameters 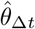 from datasets which were subsampled using different ∆*t*. The parameters were inferred by fitting up to order three moments of the data. The parameters found by the Linear MBI method were used as initial condition for the Non-linear MBI for time intervals below 0.8 min. This methods was not computationally feasible for time intervals above 0.8 min, thus we used the parameters inferred by the Non-linear MBI at time interval 0.4 min as an initial condition for those cases. The error function was then computed for every time interval ∆*t* and order *l* moment as follows,

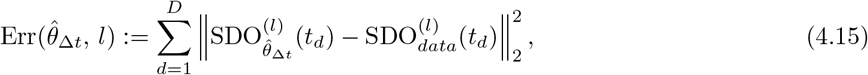

where 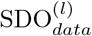 is the vector containing order *l* ∈ {1, 2, 3} moments of the data, and 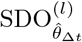 is the order *l* ∈ {1, 2, 3} moments vector, computed from 1000 SSA trajectories, which were generated with the parameters 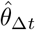 along the full dataset. We computed the error at all 130 snapshots rather than just on the subsampled snapshots in order to observe how well the approaches estimated the unseen data in between the fitted data.

#### 4.8.2 Comparison of the MBI methods

We compared the network inferred from the Linear MBI and the Non-linear MBI by using the Akaike Information Criterion (AIC). The AIC was used to rank inference models by considering a trade-off between goodness of fit and overfitting (53). To account for the small number of snapshot data points fitted in the MBI methods, which decreased from 130 to only 5 as we increased the time interval separations between snapshots, we used AIC_*c*_, which corrected the original AIC for small sample sizes (54). Is is defined as follows,

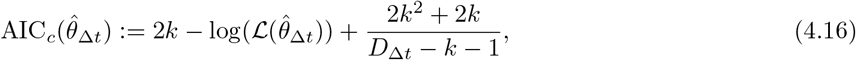

where *k* is the number of inferred parameters, 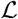 is the likelihood function, and *D*_∆*t*_ ∈ {130, 65, 33, 17, 13, 11, 9, 5} is the number of snapshot data points subsampled respectively corresponding to each ∆*t*.

For the MBI methods, the negative log-likelihood function was proportional to the sum of errors in the mean and variance. The AIC_*c*_ then reduces to,

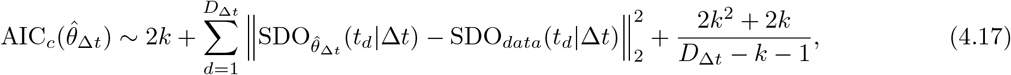

where SDO_*data*_ is the moment vector containing up to order two moments of the data, and 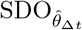 is the moment vector containing up to order two moments computed from 1000 SSA trajectories, which were generated with the parameters 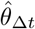 along the full dataset. The argument (*τ*_*d*_|∆*t*) is used to indicate that the SDO vectors only contain the snapshot data points subsampled with ∆*t*.

## A Supplementary Methods

### A.1 Markov Jump Process

To derive an expression of the moments, we begin by modeling the interactions within a system of *N*_*s*_ species as a stochastic process representing the number of species undergoing *N*_*r*_ reactions. This process is well described as a jump Markov process known as *Kurtz process* which describes the population count configuration 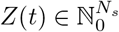 of the species at time *t* as

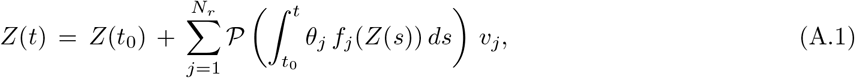

where 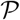 is an inhomogeneous Poisson process. *t*_0_ is initial time, *θ*_*j*_ *f*_*j*_ is the propensity function representing the rate at which the *j*-th reaction fires. And 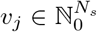 is the stoichiometry vector representing the change in species count through the *j*-th reaction.

It was shown (55) that Equation A.1 leads to the well known *Chemical Master Equation*, which represents the time evolution of the probability distribution *p* of the species count configuration *Z*(*t*) as follows

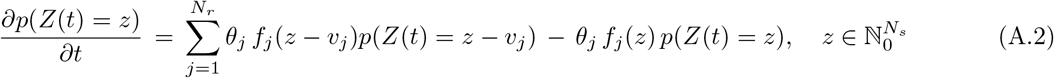

Then from Equation A.2, it can be shown (56) that for any monomial function *φ*, the derivatives of the expectation of *φ*(*Z*(*t*)) is given by

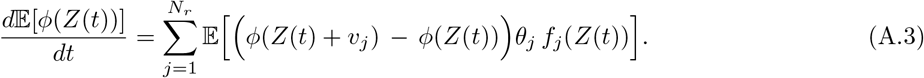

Using Equation A.3, we can write down the raw moments, *m*(*t*), of the process, *Z*(*t*), as a linear system of ODEs.

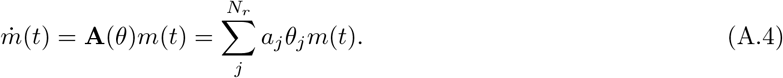

**A** is known as the *design matrix* and is a linear matrix in the propensitie coefficients 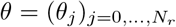, and *a*_*j*_ refers to the *design block* of **A** corresponding to the stoichiometry 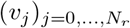 of each reaction *j* that we allow the GRN to undergo.

### A.2 SINDy

Sparse Identification of Nonlinear Dynamics (SINDy) was develop to discover the nonlinear dynamical systems that accurately represents the data (57). Given the dynamical system

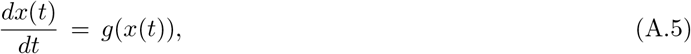

where *x*(*t*) ∈ ℝ^*S*^ is the vector of dimension *S* ∈ ℕ representing the state of the system at time *t*. SINDy method discovers the function *g* by using *D* snapshot data points (measured at time *t*_0_, … . *t*_*D*_) and by applying sparse regression techniques. The first step is to generate a library of *K* candidate nonlinear functions of snapshot data **X** = (*x*(*t*_1_), … . *x*(*t*_D_))^*T*^ ∈ ℝ^*D*×*S*^ as

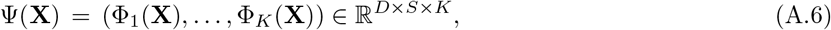

where

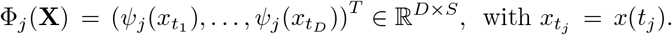

The next step is to formulate the linear system,

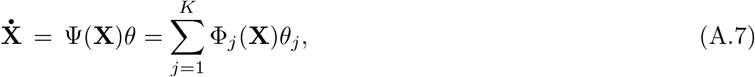

where *θ* = (*θ*_1_, … . *θ*_*K*_)^*T*^ are is vector consisting of the parameters of the nonlinearities that we consider.

The derivatives 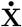 are numerically approximated from the data. The last step is to apply sparse regression techniques to find the sparse vectors parameters 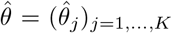 reflecting which nonlinearities are occurring in the system in Equation A.7. We can then denote 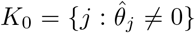 and reconstruct the nonlinear function *g* in Eq. A.5 as follows,

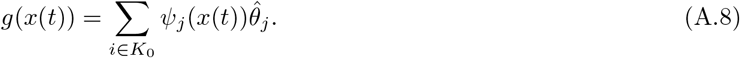

## B Supplementary Results

### B.1 GRN inference is sensitive to starting populations

The under-performance of the inference methods on the Mono-I model data was surprising. To investigate if the direction from which the target state is approached could influence the inference of the correct regulatory direction and mechanism, we simulated the Mono-I model with different starting population counts of (mRNA A, mRNA B): (70, 0), (0, 70) and (70, 70) (Fig. D Top).

Firstly, we found that all methods struggled in accurately capturing the right GRN, each method failing at different starting points. SINCERITIES was able to detect that there was only one interaction in all scenarios, however, it only captured the right GRN for starting population (70, 0). That is, when the starting population had an abundance of mRNA A, it could detect that A was being repressed (Fig. D a). Interestingly, SINCERITIES predicted the right GRN in all the replicates for the case (70, 0) (Supp. Fig. B a) The Linear MBI method did not predict the right GRN for any of the starting populations. At best, for population (70, 70), Linear MBI predicted the right GRN 26 % of the time (Supp. Fig. B b). Surprisingly, in non-symmetric starting population cases, it did not capture some of the self-regulation edges (Fig. D b). Lastly, the Nonlinear MBI under-performed when we started with an abundance of mRNA B. Looking into the replicates, we found that it predicted the right GRN 20 % of the time in the (0, 70) case and 40 % of the time in the (70, 70) case (Fig. D c, Supp. Fig. B c).

In summary, we saw that the direction from which we approach the target state influences the outcome of the inference of current methods, and this “directional bias” needs to be considered.

## Supplementary Figures

**Supplementary Figure A:**
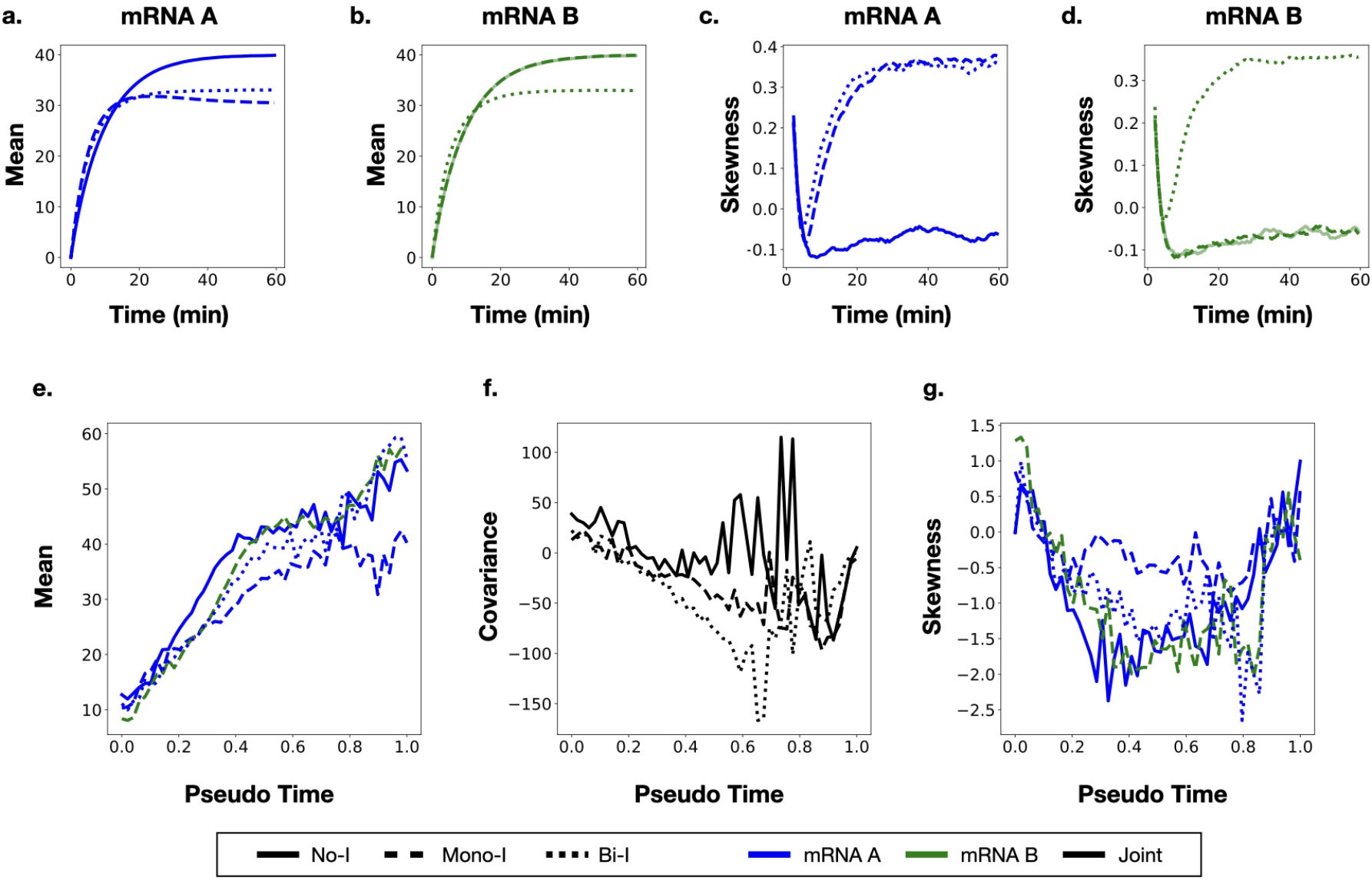
Comparison of the moments/statistics between simulated data (top row) and Pseudo Time augmented data (bottom row) for the three two-gene interaction models No-I, Mono-I, and Bi-I.

**Supplementary Figure B:**
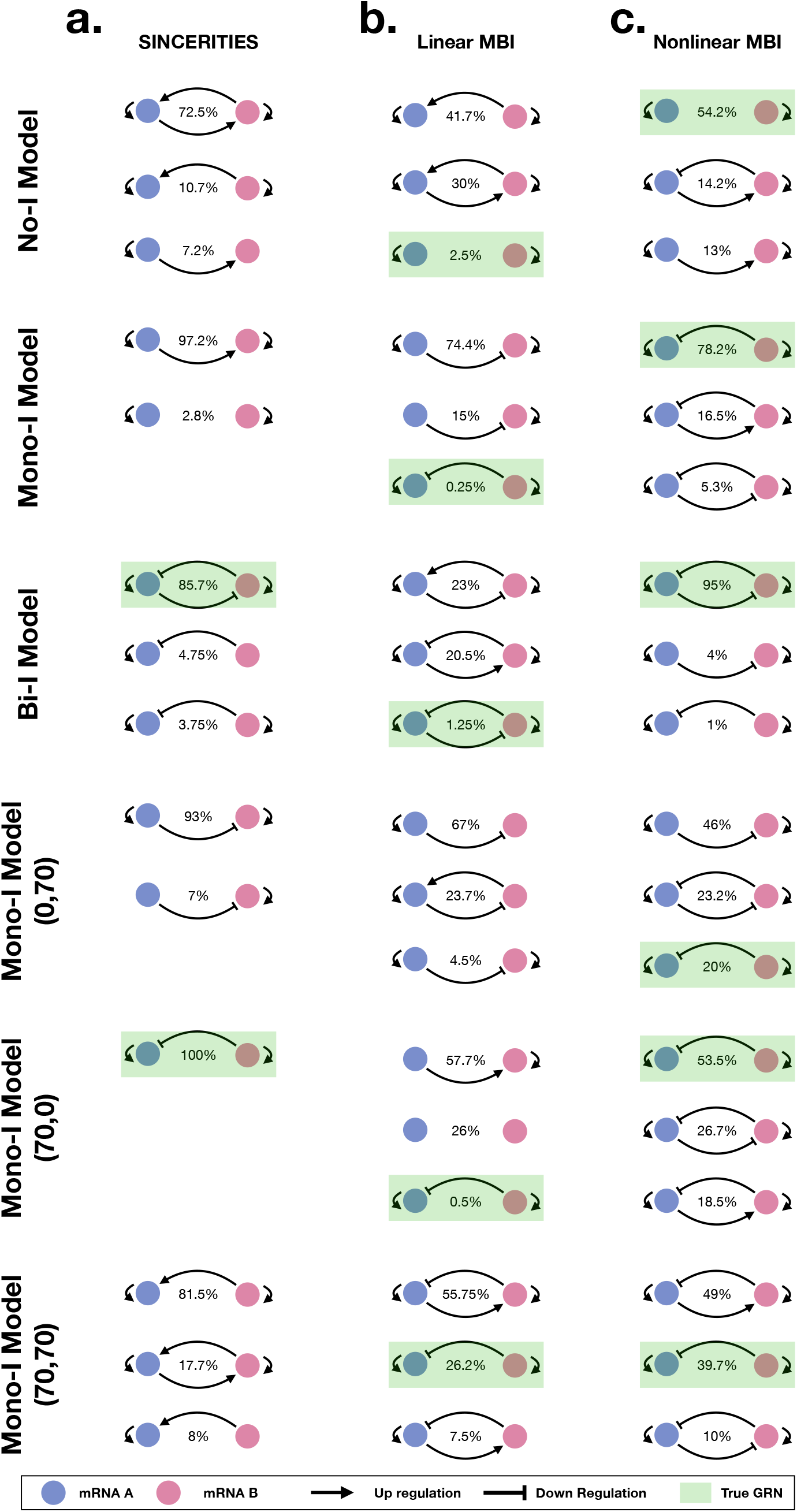
Predicted GRNs and the percentage of times they were inferred by the methods within a batch of 400 snapshot time series replicates datasets (Section 4.3): **a.** SINCERITIES, **b.** Linear MBI, **c.** Nonlinear MBI.

**Supplementary Figure C:**
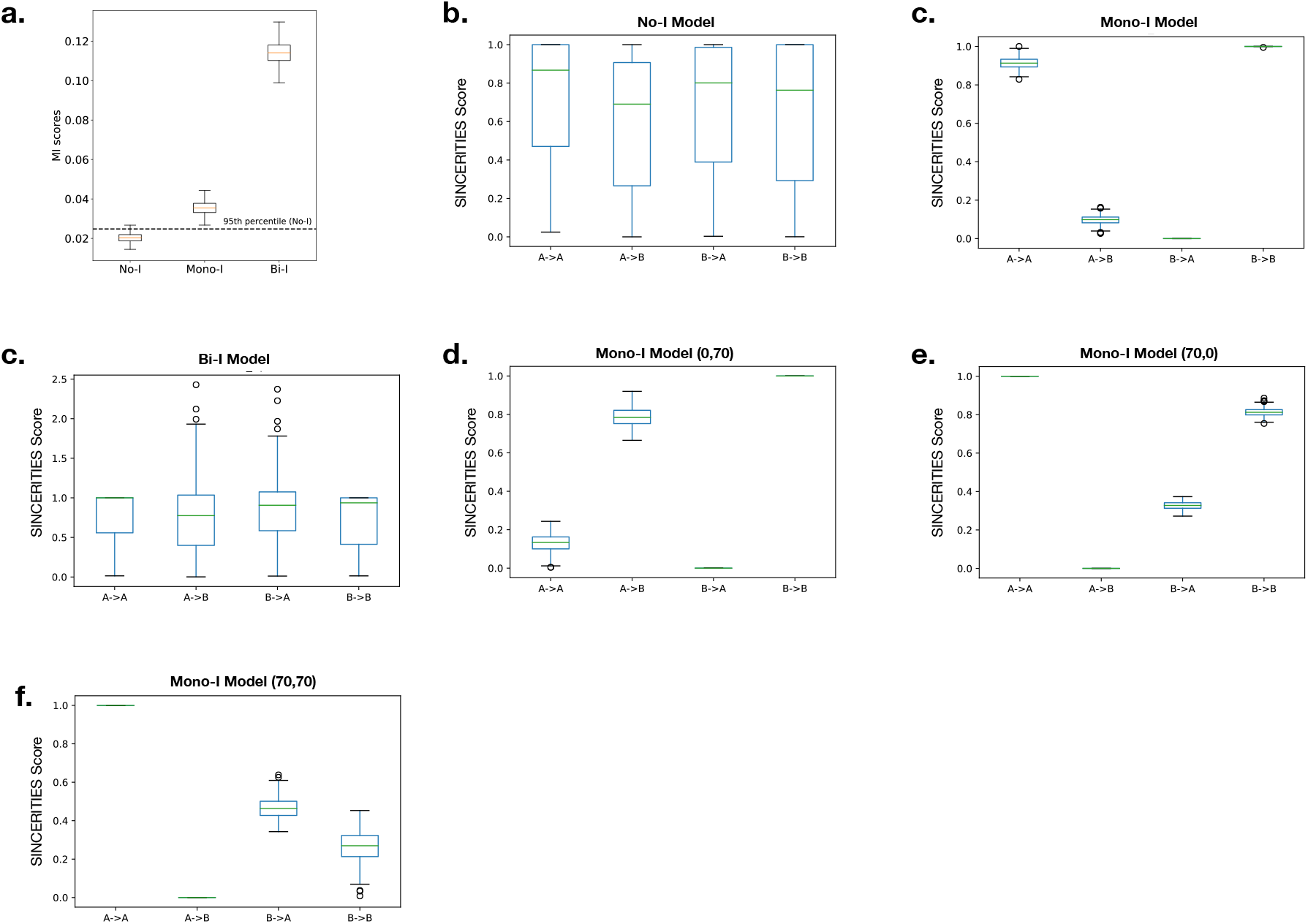
Comparison of interaction scores for the three GRN models: No-I, Mono-I, and Bi-I: (**a.**) MI scores with initial population counts (0, 0), (**b., c., d.**) SINCERITIES scores from data simulated with initial mRNA population counts (0, 0), (**e., f., g.**) SINCERITIES scores for the data simulated with initial mRNA population counts (0,70), (70, 0), and (70, 70).

**Supplementary Figure D:**
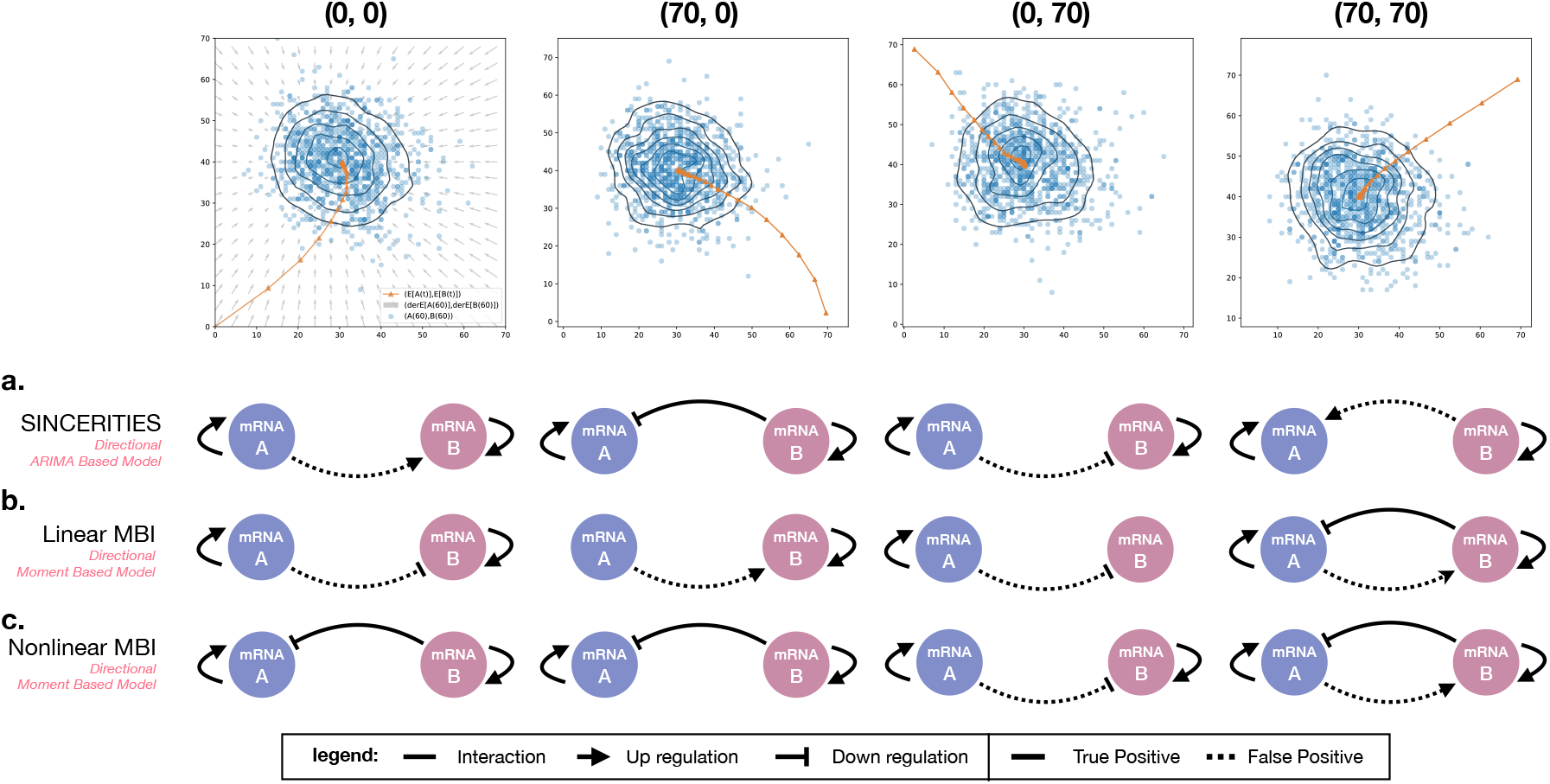
GRNs for different starting mRNA count configurations of the Mono-I model. Illustration of GRNs obtained for data generated from the Mono-I model with different starting population count configurations for (mRNA A, mRNA B): (0, 0), (70, 0), (0, 70), and (70, 70). Top row: 1000 sample mRNA population counts at time *T* = 60, vector fields indicate the direction of change in the model using derivatives of the first order moments at a given point in the plane, and the orange line is the mean expression trajectory in the data. The three rows **a., b.,** and **c.** show the inferred GRNs aligned with the corresponding initial conditions in the top row (refer to Fig. 4 for the meaning of the graphs).

**Supplementary Figure E:**
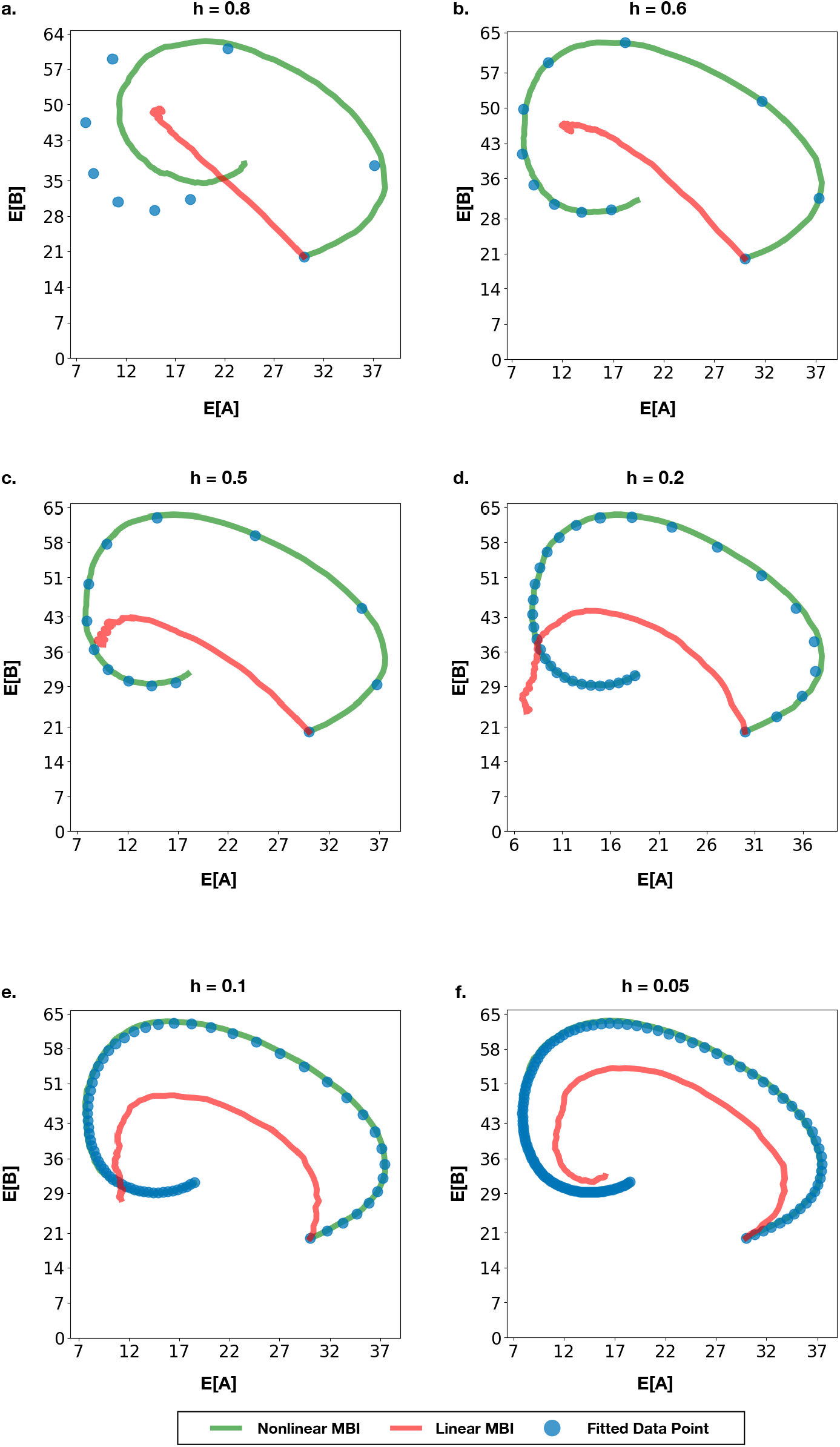
Comparison of the first moments time course reconstruction of the stochastic damped oscillator model for different snapshots time-series data subsampled at time intervals h (min) between snapshots. Time course reconstruction were generated from SSA of the model using the parameters inferred by each MBI methods.

## References

[1] Stegle, O., Teichmann, S. A. & Marioni, J. C. Computational and analytical challenges in single-cell transcriptomics. Nature Reviews Genetics 16, 133–145 (2015).

[2] Kolodziejczyk, A. A., Kim, J. K., Svensson, V., Marioni, J. C. & Teichmann, S. A. The technology and biology of single-cell RNA sequencing. Molecular Cell 58, 610–620 (2015).

[3] Fiers, M. W. E. J. et al. Mapping gene regulatory networks from single-cell omics data. Briefings in Functional Genomics 17, 246–254 (2018).

[4] Ghanbari, M., Lasserre, J. & Vingron, M. The distance precision matrix: computing networks from non-linear relationships. Bioinformatics 35, 1009–1017 (2018).

[5] Hwang, B., Lee, J. H. & Bang, D. Single-cell RNA sequencing technologies and bioinformatics pipelines. Experimental 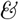 Molecular Medicine 50 (2018).

[6] Giovanni, I., Ramon, M.-B. & Holger, H. Single-cell transcriptomics unveils gene regulatory network plasticity. Genome Biology 20 (2019).

[7] Delgado, F. M. & Gómez-Vela, F. Computational methods for gene regulatory networks reconstruction and analysis: A review. Artificial Intelligence in Medicine 95, 133–145 (2019).

[8] Trapnell, C. et al. The dynamics and regulators of cell fate decisions are revealed by pseudotemporal ordering of single cells. Nature Biotechnology 32, 381–386 (2014).

[9] Qiu, X. et al. Reversed graph embedding resolves complex single-cell trajectories. Nature Methods 14, 979–982 (2017).

[10] La Manno, G. et al. Rna velocity of single cells. Nature 560, 494–498 (2018).

[11] Pratapa, A., Jalihal, A. P., Law, J. N., Bharadwaj, A. & Murali, T. M. Benchmarking algorithms for gene regulatory network inference from single-cell transcriptomic data. Nature Methods 17, 147–154 (2020).

[12] Kitano, H. Systems biology: A brief overview. Science 295, 1662–1664 (2002).

[13] Enze, L., Lang, L. & Lijun, C. Gene regulatory network review. In Encyclopedia of Bioinformatics and Computational Biology, 155–164 (Elsevier, 2019).

[14] Holehouse, J., Cao, Z. & Grima, R. Stochastic modeling of autoregulatory genetic feedback loops: A review and comparative study. Biophysical Journal 118, 1517 – 1525 (2020).

[15] Davidson, E. H. et al. A provisional regulatory gene network for specification of endomesoderm in the sea urchin embryo. Developmental Biology 246, 162–190 (2002).

[16] Streit, A. et al. Experimental approaches for gene regulatory network construction: The chick as a model system. Genesis 51, 296–310 (2013).

[17] Zheng, G. & Huang, T. The reconstruction and analysis of gene regulatory networks. In Methods in Molecular Biology, 137–154 (Springer New York, 2018).

[18] Barbuti, R., Gori, R., Milazzo, P. & Nasti, L. A survey of gene regulatory networks modelling methods: from differential equations, to boolean and qualitative bioinspired models. Journal of Membrane Computing (2020).

[19] Vogel, C. & Marcotte, E. M. Insights into the regulation of protein abundance from proteomic and transcriptomic analyses. Nature Reviews Genetics 13, 227–232 (2012).

[20] Fortelny, N., Overall, C. M., Pavlidis, P. & Freue, G. V. C. Can we predict protein from mRNA levels? Nature 547, E19–E20 (2017).

[21] S., K. ppcor: An r package for a fast calculation to semi-partial correlation coefficients. Commun. Stat. Appl. Methods 22, 665–674 (2015).

[22] Chan, T. E., Stumpf, M. P. H. & Babtie, A. C. Gene regulatory network inference from single-cell data using multivariate information measures. Cell Systems 5, 251–267.e3 (2017).

[23] Spetch, A. T. & Li, J. Leap: constructing gene co-expression networks for single-cell rna-sequencing data using pseudotime ordering. Bioinformatics 33, 764–766 (2017).

[24] Papili Gao, N., Ud-Dean, S. M. M., Gandrillon, O. & Gunawan, R. SINCERITIES: inferring gene regulatory networks from time-stamped single cell transcriptional expression profiles. Bioinformatics 34, 258–266 (2018).

[25] Bonnaffoux, A. et al. WASABI: a dynamic iterative framework for gene regulatory network inference. BMC Bioinformatics 20 (2019).

[26] Klimovskaia, A., Ganscha, S. & Claassen, M. Sparse regression based structure learning of stochastic reaction networks from single cell snapshot time series. PLOS Computational Biology 12, 1–20 (2016).

[27] Matsumoto, H. et al. SCODE: an efficient regulatory network inference algorithm from single-cell RNA-seq during differentiation. Bioinformatics 33, 2314–2321 (2017).

[28] Aubin Frankowski, P. C. & Vert, J. P. Gene regulation inference from single-cell RNA-seq data with linear differential equations and velocity inference. Bioinformatics (2020). Btaa576.

[29] Haghverdi, L., Büttner, M., Wolf, F. A., Buettner, F. & Theis, F. J. Diffusion pseudotime robustly reconstructs lineage branching. Nature Methods 13, 845–848 (2016).

[30] Eraslan, G., Simon, L. M., Mircea, M., Mueller, N. S. & Theis, F. J. Single-cell rna-seq denoising using a deep count autoencoder. Nature communications 10, 390–390 (2019).

[31] Cao, Z. & Grima, R. Analytical distributions for detailed models of stochastic gene expression in eukaryotic cells. Proceedings of the National Academy of Sciences 117, 4682–4692 (2020).

[32] Ko, M. S. A stochastic model for gene induction. Journal of Theoretical Biology 153, 181 –194 (1991).

[33] McAdams, H. H. & Arkin, A. Stochastic mechanisms in gene expression. Proceedings of the National Academy of Sciences 94, 814–819 (1997).

[34] Thattai, M. & van Oudenaarden, A. Intrinsic noise in gene regulatory networks. Proceedings of the National Academy of Sciences of the United States of America 98, 8614–8619 (2001).

[35] Swain, P. S., Elowitz, M. B. & Siggia, E. D. Intrinsic and extrinsic contributions to stochasticity in gene expression. Proceedings of the National Academy of Sciences 99, 12795–12800 (2002).

[36] Dibaeinia, P. & Sinha, S. Sergio: A single-cell expression simulator guided by gene regulatory networks. Cell Systems 11, 252 – 271.e11 (2020).

[37] LĹ’ahnemann, D. & al. Eleven grand challenges in single-cell data science. Genome Biology 21 (2020).

[38] Tanay, A. & Regev, A. Scaling single-cell genomics from phenomenology to mechanism. Nature 541 (2017).

[39] Padi, M. & Quackenbush, J. Integrating transcriptional and protein interaction networks to prioritize condition-specific master regulators. BMC Systems Biology 9, 80 (2015).

[40] Chu, L.-F. et al. Single-cell RNA-seq reveals novel regulators of human embryonic stem cell differentiation to definitive endoderm. Genome Biology 17 (2016).

[41] Kouno, T. et al. Temporal dynamics and transcriptional control using single-cell gene expression analysis. Genome Biology 14 (2013).

[42] Stumpf, P. S. et al. Stem cell differentiation as a non-markov stochastic process. Cell Systems 5, 268–282.e7 (2017).

[43] Marbach, D., Schaffter, T., Mattiussi, C. & Floreano, D. Generating realistic in silico gene networks for performance assessment of reverse engineering methods. J Comput Biol 16, 229–239 (2009).

[44] Marbach, D. et al. Revealing strengths and weaknesses of methods for gene network inference. Proceedings of the National Academy of Sciences 107, 6286–6291 (2010).

[45] Gillespie, D. T. A General method for numerically simulating the stochastic time evolution of coupled chemical reactions. Journal of Computational Physics 22, 403–434 (1976).

[46] Wolf, F. A., Angerer, P. & Theis, F. J. SCANPY: large-scale single-cell gene expression data analysis. Genome Biology 19 (2018).

[47] Magwene, P. M., Lizardi, P. & Kim, J. Reconstructing the temporal ordering of biological samples using microarray data. Bioinformatics 19, 842–850 (2003).

[48] Pierre-Cyril, A.-F. & Jean-Philippe, V. Gene regulation inference from single-cell RNA-seq data with linear differential equations and velocity inference. Bioinformatics (2020).

[49] Hoffmann, M., Fröohner, C. & Noée, F. Reactive SINDy: Discovering governing reactions from concentration data. The Journal of Chemical Physics 150, 025101 (2019).

[50] Leclerc, R. D. Survival of the sparsest: robust gene networks are parsimonious. Molecular Systems Biology 4 (2008).

[51] Gaines, B. R., Kim, J. & Zhou, H. Algorithms for fitting the constrained lasso. Journal of Computational and Graphical Statistics 27, 861–871 (2018).

[52] Virtanen, P. et al. SciPy 1.0: fundamental algorithms for scientific computing in python. Nature Methods 17, 261–272 (2020).

[53] Akaike, H. A new look at the statistical model identification. IEEE Transactions on Automatic Control 19, 716–723 (1974).

[54] Burnham, K. P. & Anderson, D. R. Multimodel inference: Understanding aic and bic in model selection. Sociological Methods 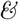 Research 33, 261–304 (2004).

[55] Sunkara, V. Algebraic expressions of conditional expectations in gene regulatory networks. Journal of Mathematical Biology 79, 1779–1829 (2019).

[56] Engblom, S. Computing the moments of high dimensional solutions of the master equation. Applied Mathematics and Computation 180, 498 – 515 (2006).

[57] Brunton, S. L., Proctor, J. L. & Kutz, J. N. Discovering governing equations from data by sparse identification of nonlinear dynamical systems. Proceedings of the National Academy of Sciences 113, 3932–3937 (2016).

